# One-shot entorhinal maps enable flexible navigation in novel environments

**DOI:** 10.1101/2023.09.07.556744

**Authors:** John H. Wen, Ben Sorscher, Surya Ganguli, Lisa M Giocomo

## Abstract

Animals face the substantial challenge of navigating novel environments to find food, shelter, or mates. In mammals, hexagonal grid cells in the medial entorhinal cortex create a map-like population representation of the external environment ^1–7^. However, it remains unclear how the grid map can adapt to novel environmental features on a rapid, behaviorally relevant time scale. By recording over fifteen thousand grid cells in mice navigating virtual environments, we found grid cell activity was weakly anchored to landmark inputs through a *fixed* circuit relationship. A computational model based on this fixed circuit assumption accurately predicted grid spatial patterns in environments with novel landmark rearrangements. Finally, a medial entorhinal cortex-dependent task revealed that while grid cell firing patterns remain anchored to landmarks, behavior can adapt to changes in landmark location via a downstream region implementing behavioral time scale synaptic plasticity ^8^. This fixed but weak anchoring of grid cells to landmarks endows the grid map with powerful computational properties. The fixed nature allows the generation of rapid stable maps for novel environments after a *single* exposure. The weak nature allows these rapidly formed maps to incur only small distortions between distances traveled in real versus neural space. Overall, such rapid low distortion mapping can then mediate accurate navigational behavior in rapidly changing environments through downstream plasticity.

## Main Text

To navigate to an intended location, the brain builds a map of space that integrates external landmark information with idiothetic cues derived from self-motion ^1,9–12^. A challenge faced by animals exploring naturalistic environments is the near constant need to incorporate dynamically changing environmental features into their existing maps of space. Such changes can be caused by weather patterns, natural forces, or competition for resources, occurring over long (e.g., seasonal changes) and short (e.g., destruction of a nest site) timescales. How the brain rapidly incorporates such novel features into an animal’s internal map of space to support accurate navigation remains incompletely understood.

One potential solution to this challenge is a rigid system for calculating distance traveled through space using only idiothetic cues, such as path integration ^13–15^. Such a system can operate instantly in any environment and calculate distance traveled regardless of changes in environmental features. However, path integration systems untethered to landmarks are susceptible to noise and thus accumulate error ^16,17^. In contrast, a slower but less error-prone system is one where internal maps of space integrate novel environmental features, such as a change in the location of a visual landmark, via Hebbian synaptic mechanisms ^18–20^. This solution allows the neural map of space to flexibly reorganize to accommodate changes in the environment, enabling faithful internal maps of the external world. That is, distances and directions traveled in real space would be in correspondence to ones represented by trajectories in neural space ^21^. However, it comes with the disadvantage that Hebbian learning takes time and repeated experience. Even so, studies in *Drosophila* provide strong evidence consistent with this mechanism: Hebbian plasticity operates on the timescale of minutes to integrate the location of novel landmarks in the fly’s internal map of heading orientation ^18,19^.

While studies in *Drosophila* have pointed to Hebbian plasticity to integrate novel landmarks into existing internal maps, how the mammalian spatial mapping system accomplishes this is unknown. Interestingly, neither solution seems to be entirely consistent with previous works on the medial entorhinal cortex (MEC) grid cell system, which is proposed to provide a neural substrate for building an internal map of the external world for navigation ^1,9,22–24^. While grid cells use idiothetic cues to encode distance traveled in the dark ^25,26^, the spatial regularity of their firing patterns is sensitive to changes in environmental features. For example, boundaries ^27–31^ and locations associated with reward ^32,33^ can fracture and distort grid cell firing patterns, while also providing an error-correcting mechanism for internal position estimates ^29^. Such sensitivity to environmental features rules out the first solution of a highly rigid system that supports navigation only based on idiothetic cues. Additionally, geometric perturbations to familiar environments ^30,34,35^ and exposure to novel geometries ^36,37^ result in distortions in grid cell firing patterns that can persist for days ^34–13;36,38^. These persistent distortions in response to environmental changes appear to be inconsistent with the rapid Hebbian plasticity solution observed in *Drosophila*. On a longer timescale, over many days, grid cells can eventually generate a coherent map for compartmentalized environments ^28,39^. However, such long timescale plasticity cannot account for how animals rapidly and efficiently map novel spaces for navigation ^40^, nor can it account for short timescale changes to naturalistic environments.

Here, we asked whether a third possibility might exist to explain how mammals rapidly map novel landmark rearrangements or entirely novel environments. To address this mystery, we used Neuropixels probes to record from more than fifteen thousand MEC grid cells as mice navigated virtual reality environments in which novel and familiar landmarks were systematically manipulated. Large numbers of simultaneously recorded grid cells allowed us to consider how grid firing patterns, referred to here as the “grid map”, rapidly incorporated novel environmental features in the context of attractor networks, a computational framework highly consistent with the emergence of grid cell firing patterns ^9,41–50^. Altogether, our results point to grid cells as incorporating landmark information through the dynamics of a *fixed* circuit, thereby endowing the grid cell system with the powerful computational capacity of one-shot mapping of novel environments without any need for plasticity. However, this rapid one-shot mapping comes at the cost of distortions in the grid map. Nevertheless, despite their susceptibility to landmark-induced distortions, the rapidly forming, stable grid maps appeared to be both necessary and sufficient to solve a behavioral task that required integrating self-motion with external landmarks.

### Grid cell attractor dynamics persist in the absence of visual features

To investigate how MEC grid cells respond to novel environments, we used Neuropixels probes ^51^ to record from tens of thousands of MEC neurons (n = 68,484 neurons, Extended Data Fig. 1, Extended Data Table 1) in head-fixed mice navigating virtual reality (VR) 1-dimensional (1D) linear tracks. For all experiments, mice ran a block of trials in complete darkness (Methods). In the dark, single-cell spike trains exhibited little apparent spatial structure (Fig. 1a). However, applying the Fourier transform to each neuron’s spatially binned spike train revealed a distinctive three-peaked structure in a subset of recorded neurons, which we classified as grid cells for several reasons (n = 15,342, Methods) (Fig. 1b). First, this three-peaked structure suggested that over short distance scales (∼10 laps), the activity of grid cells in 1D VR was well-described by a 1D slice through a 2D hexagonal lattice, consistent with prior work ^52^. Second, grid cells clustered into discrete modules by their power spectra (Fig. 1c) and, in sessions with multiple modules, the inferred distance between grid firing nodes (i.e. grid scale) increased in discrete steps with a scale ratio between 1.4 and 1.7 (Fig. 1d), cardinal features of grid cells ^30^. Third, grid cells from the same module preserved their correlation structures and phase relationships across long running distances (Extended Data Fig. 2) ^41,43,45^.

**Figure 1.**
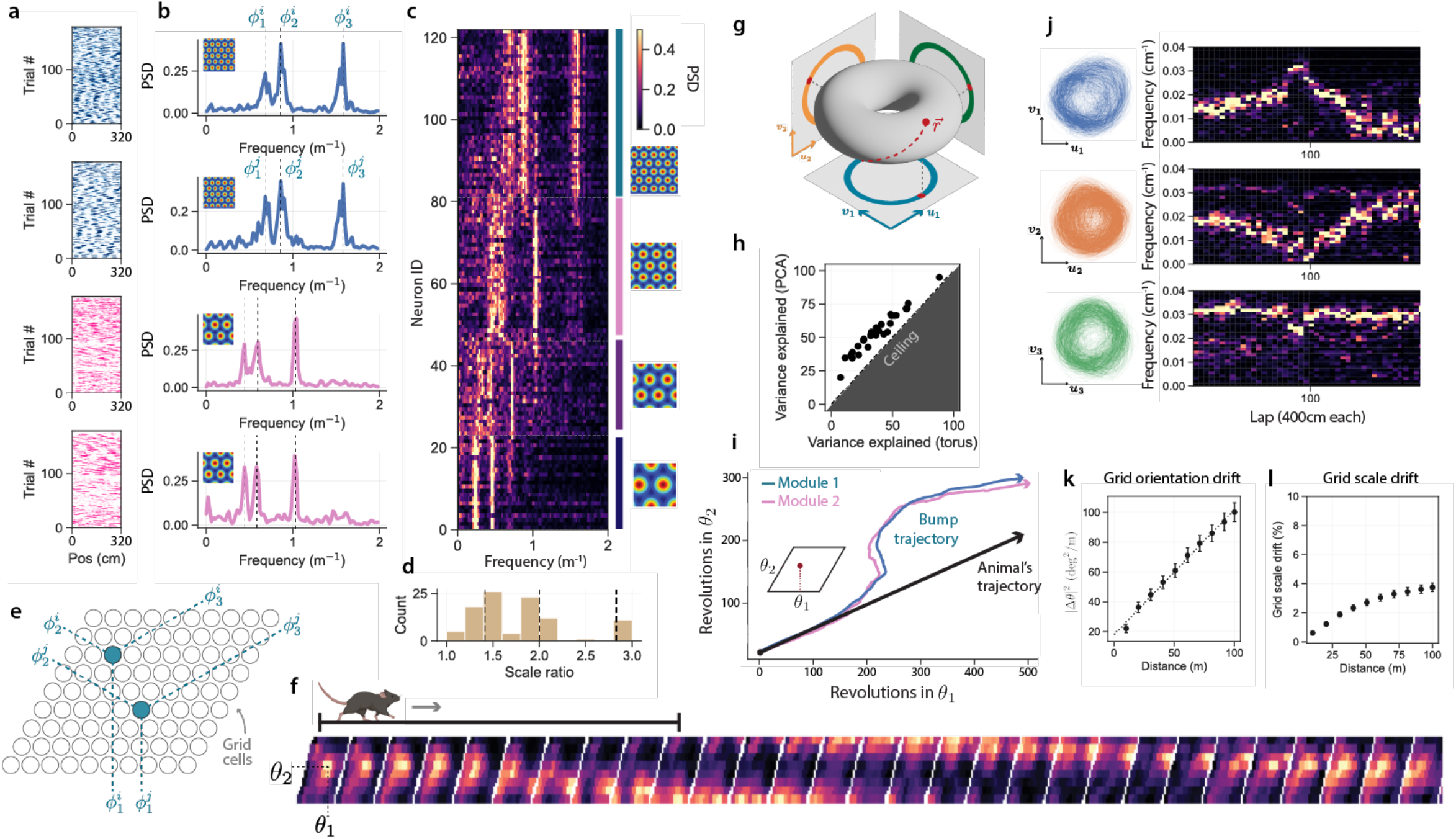
The activity of MEC neurons is well described by attractor dynamics in the dark. **a**. Example spike raster plots for four grid cells from two simultaneously recorded modules over multiple trials in the dark. Dots indicate spikes, color coded to match the modules in (c). One lap equals 320 cm distance traveled. **b**. Power spectral densities (PSDs) for each cell in (a), computed over a window size of 10 laps. **c**. Power spectra for all simultaneously recorded MEC neurons in a session clustered into modules of increasing frequency. *Left*: Power spectra for each neuron color coded for maximum (white) and minimum (black) power in each frequency, sorted dorsal (top) to ventral (bottom). *Right*: Each colored bar (teal, pink, purple, dark blue) indicates a single module, illustrated by the inferred grid scale in a 2D open field (color coded for minimum (blue) and maximum (red) firing rate). **d**. Histogram of inferred 2D grid scale ratios between modules for all recorded sessions (animal n = 32, session n = 214, module n = 195, cell n = 7821). Dashed vertical lines indicate powers of 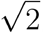. **e**. Schematic of sorting neurons onto a neural sheet. For a given module, each grid cell is sorted based on its 2D phase *ϕ*_1_ · *ϕ*_2_, as denoted in (b) (Methods). **f**. When sorted in this fashion, grid cell population activity reveals a bump of activity that translates with the animal’s forward movement in the dark. Each frame indicates the population activity of 64 co-modular grid cells within a 2 cm bin along the track, color coded for maximum (white) and minimum (black) firing rates, averaged over 20 trials. **g**. Neural trajectory (red dashed line) in high-dimensional firing rate space lies close to a torus, identified by projecting neural activity onto the three pairs of axes *u*_1,2,3_ = cos(*ϕ*_1,2,3_) and *υ*_1,2,3_ = sin(*ϕ*_1,2,3_) . **h**. The 6D subspace in (g) captures nearly as much variance in the grid cell population activity as the top 6 principal components, across modules and sessions (black dots). **i**. Inferred (unwrapped) bump trajectory for grid cells from two simultaneously recorded modules: blue module (37 grid cells) and pink module (36 grid cells). Bump trajectories deviated from a straight-line trajectory, but the modules drifted together. **j**. Population activity of a set of co-modular grid cells projected onto the three pairs of axes identified in (g). Activity lies close to a ring in each 2D subspace. The three peaks in the PSD (b) are the frequencies at which the neural activity wraps around each of these rings. These frequencies drifted over long-distance scales (*right*), color coded for maximum (white) and minimum (black) power. **k**. Slice angle drifts over 100 meters in the dark and is well approximated by a diffusion process with diffusion constant D = 1.16 ± 0.07 deg^2^/m (animal n = 4, session n = 6, module n = 10, cell n = 260). Dots indicate mean, bars indicate standard error of the mean (SEM). **l**. Grid scale remained approximately fixed over 100 meters in the dark (animal n = 4, session n = 6, module n = 10, cell n = 260). Dots indicate mean, bars indicate SEM.

**Figure 2.**
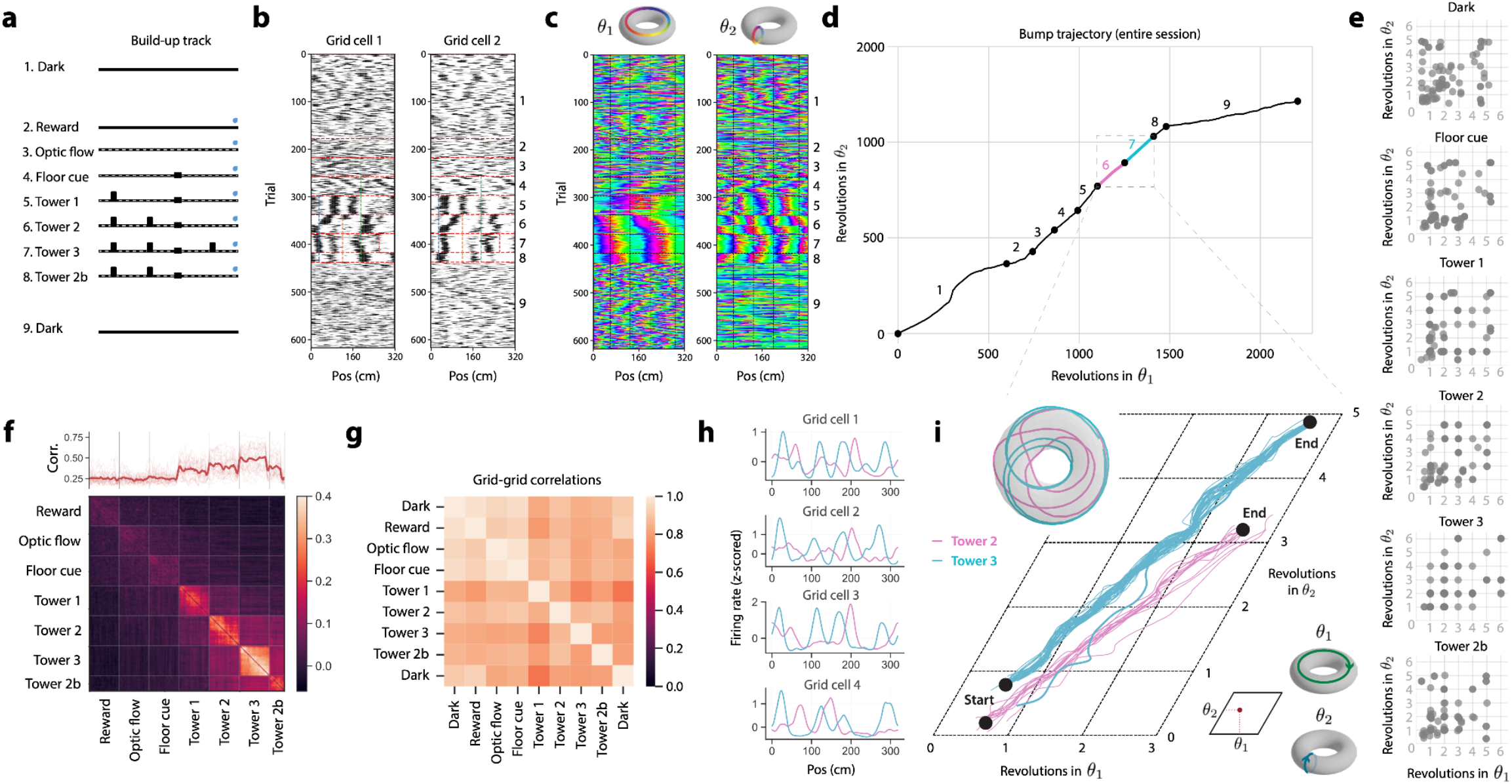
Visual landmarks entrained grid cells to periodic trajectories along the torus axes. **a**. Side view schematic of the build-up track. At the end of 320 cm, the animal teleported seamlessly to the beginning of the track. **b**. Example raster plots of two grid cells recorded from one animal (mouse identity [ID]: N2) over the one build-up track session. Black dots indicate spikes, red dotted lines indicate trial block transitions. Trial block numbers, as in (a), listed on the far right **c**. Bump trajectory for one recording session on the build-up track plotted as 2D coordinates on the torus: *ϕ*_1_ (*left*) and *ϕ*_2_ (*right*), for the same session as in (b) (cell n = 37). Colors indicate angle along each axis of the torus. **d**. Activity bump trajectory (unwrapped) for an example session (cell n = 37). Black dots indicate the start and end of the session, and addition or removal of a feature to the track. Numbers correspond to trial blocks in (a). **e**. Landmarks entrain activity bump to periodic trajectories across grid modules and sessions (each dot represents one module from one session). For each block of trials, as in (a), dots indicate the number of revolutions of the bump of activity along each axis of the torus as the animal completes one lap down the track. As landmarks are added, bump trajectories snap to whole numbers (gray lines) of revolutions around each axis of the torus (animal n = 4, session n = 14, module n = 35, cell n = 747). **f**. Grid cell activity rapidly stabilizes within each trial block but remaps between trial blocks. *Top*: moving-window correlation (Pearson r) of grid cell population activity (window size = 20 meters, thick line = mean, thin lines = individual blocks, session n = 14, animal n = 4, module n = 35, cell n = 747). *Bottom*: correlation (Pearson r) of grid cells across all trials (session n = 14, animal n = 4, module n = 35, cell n = 747). **g**. Grid-grid correlations are preserved across VR environments. Pairwise temporal correlation matrices are computed for grid cell spike-trains within each block, for the session in (b-d). Color indicates correlation (Pearson r) between correlation matrices in two blocks (see also Extended Data Fig. 2). **h**. Example tuning curves for four grid cells recorded in the example session in (b) for the 2 tower (pink) and 3 tower (blue) trial blocks (blocks 6 and 7). **i**. Example session from (d, h) on the build-up track with single-trial bump trajectories (unwrapped) for the 2 tower (pink) and 3 tower (blue) trial blocks. Traces correspond to bump trajectories on individual trials. Bold blue line (overlapping with the pink lines) indicates the first trial in which there were 3 towers. Average trajectories of the bump for each trial block are shown on the torus (*inset*).

Next, to build a framework for examining the population dynamics of grid cells in novel environments, we sorted grid cells from the same module (i.e., co-modular) recorded in the dark onto a 2D toroidal neural sheet. The resulting sheet allowed us to extract the moment by moment 2D grid cell attractor state. For each module, we sorted grid cells according to each cell’s phases 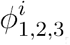, which were determined from the peaks of their power spectra (Fig. 1e) (Methods). When sorted in this manner, the grid cells’ population activity revealed a “bump” of activity that translated across the 2D toroidal neural sheet in concert with the animal’s forward movement (Fig. 1f). This is reminiscent of the activity bump observed in the *Drosophila* compass circuit that stores the fly’s internal estimate of head direction ^53^. To visualize the activity bump’s trajectory over the course of the entire experiment, we computed the 2D center of mass (COM) of the bump at each spatial bin along the track (320 cm track binned into 2 cm bins). Note that the sorting of co-modular grid cells onto the 2D toroidal neural sheet was based on the first 10 trials of the session, which allowed us to follow the subsequent trajectory of the activity bump over the course of the recording session.

Extracting the activity bump’s COM is mathematically equivalent to projecting the high-dimensional neural population activity onto a 6-dimensional subspace spanned by the vectors 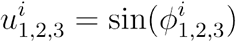 and 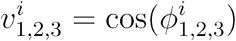, where *i* = 1, …,*N* indexes grid cells within a module (Fig. 1g). When projected onto each pair of axes, the neural activity traced out a ring (Fig. 1g). This indicated that, even in 1D and in the dark, the grid cell population activity lay close to a manifold with the topology of a twisted torus, akin to that observed in rodents navigating 2D environments with visual cues ^48^. Importantly, this torus does not live in a carefully-chosen subspace that contains very little of the total variance of the population activity. Rather, across modules and recordings, this 6D subspace explained nearly all the variance achievable by any 6D subspace (Fig. 1h). Together, these results indicate that even in the absence of visual landmarks, the grid cell population is well-described by low-dimensional attractor dynamics.

Finally, we considered the activity bump’s COM over long distances traveled in the dark. We observed that the bump’s trajectory deviated from a straight-line path (Fig. 1i). As a consequence, the 1D slice angle through the 2D lattice, parameterized by the locations of the three peaks (*f*_1_ · *f*_2_, *f*_3_) in the Fourier spectrum (Fig. 1j), drifted and rotated (Fig. 1k). For sessions in which a sufficient number of co-modular grid cells were recorded (Methods), we found the slice angle drift grew as the square root of the distance traveled, closely matching an angular diffusion process with diffusion constant D = 1.16 ± 0.07 deg^2^/m (animal n = 4, session n = 6, module n = 10, cell n = 260). The overall grid scale, on the other hand, remained stable and drifted less than 5% over 100 meters of running (Fig. 1l) (animal n = 4, session n = 6, module n = 10, cell n = 260). In recordings that captured the activity of more than one module of grid cells, we observed that activity bumps from different modules were tightly coupled and drifted in concert, taking the same overall trajectories (Fig. 1i, Extended Data Fig. 3) ^45^. These observations are consistent with the rapid accumulation of error observed in path integration estimates in the absence of sensory landmark cues ^26,29,54–56^.

**Figure 3.**
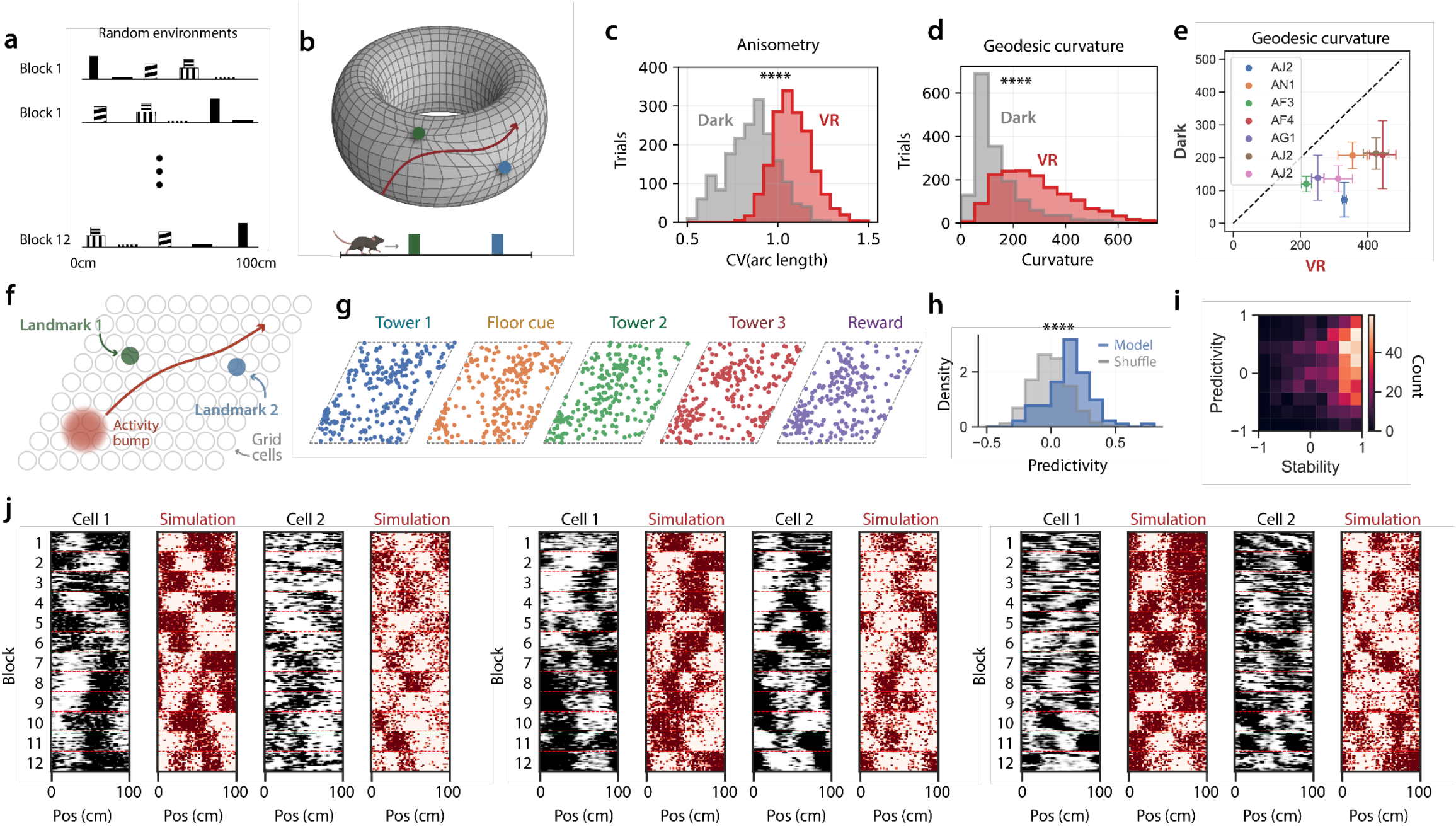
Dynamical structure underlying grid cell remapping. **a**. Schematic of the random environment track. **b**. Schematic of anisotropy and geodesic curvature (red line) for bump trajectories on the torus. Grid lines represent unit distance intervals in physical space. A perfectly isometric neural map would be characterized by regular, square grid lines. Anisometry measures the degree to which these grid lines are distorted. Similarly, a neural trajectory (red line) faithful to the animal’s true trajectory would follow a geodesic (straight-line) trajectory on the torus. Geodesic curvature measures the degree of deviation from straight-line trajectories. **c**. Bump trajectories have significantly higher anisometry in the presence of visual landmarks compared to complete darkness (KS test, p < 1e-10, animal n = 6, session n = 7), as quantified by the coefficient of variation in the arc length of neural trajectories corresponding to 16-meter trajectories in real space (Methods). **d**. Bump trajectories have significantly higher geodesic curvature in the presence of visual landmarks compared to complete darkness (KS test, p < 1e-10, animal n = 6, session n = 7). **e**. Geodesic curvature was higher in the presence of visual landmarks compared to complete darkness for all animals. Individual animals plotted as colored dots, lines indicate SEM, averaged over sessions. **f**. Cartoon of a simplified model of the error-correcting effect of landmarks on the bump trajectory. When the animal approaches a landmark (colored as in (b)), the bump is tugged toward a preferential location (pinning phase) on the neural sheet. This pinning is strong enough to stabilize the bump’s trajectory. However, it is not so strong that the bump is pinned to the same location on the neural sheet each time the animal passes a particular landmark. **g**. Location of bump on the neural sheet each time the animal passes a landmark across all trial blocks in (a) for a single grid cell module during a single session (n = 23 modules). **h**. Histogram of model predictivity (Pearson’s r) across all cells compared to a random shuffle (see Extended Data Fig. 6 for additional ablations) (KS test, p < 1e-10, animal n = 3, session n = 3, module n = 3, cell n = 91). **i**. Heatmap of model predictivity (average Pearson r between simulated grid cell tuning curves and true grid cell tuning curves on held out blocks) and grid cell stability (average Pearson r correlation between a grid cell’s tuning curve in the first half of a block of trials and the second half) (animal n = 3, session n = 3, module n = 3, cell n = 91). **j**. Example raster plots of real (black) and simulated (red) neurons from three different recordings across three different mice (two neurons from each mouse). Individual mice are denoted on the top of each set of neurons.

### Novel landmarks entrain grid cell dynamics and induce one-shot remapping

Next, we applied our method for extracting the moment-by-moment 2D grid cell attractor state (i.e. the bump COM) to assess how grid cells incorporate novel environmental features into their spatial map. We designed a VR task in which features were systematically added to the VR track (the “build-up track”). After an initial block of dark trials, animals ran 40 trials in each of blocks 2 through 7, in which features were added cumulatively (Fig. 2a). In block 2, automatic water rewards were dispensed every 320 cm. In block 3, optic flow was added to the floor of the track. In blocks 4 to 7, distinct towers were added at fixed positions. In block 8 (20 trials), we removed one of the towers. In block 9, the animal again ran a block of dark trials. Animals did not have any prior experience with any of the visual features before the first recording session.

First, we visualized the responses of individual grid cells to the addition of landmarks (Fig. 2b). We observed that as landmarks were added, more apparent spatial structure emerged within the firing patterns of grid cells. To assess the grid cell population response to the addition of landmarks, we used the phases extracted while the animal was running in the dark (as in Fig. 1) and plotted the trajectory of the activity bump across the 2D toroidal neural sheet (Fig. 2c). In particular, plotting each coordinate on the neural sheet separately (*ϕ*_1_ or *ϕ*_2_) revealed that as landmarks were added, the activity bump was entrained to whole number revolutions around the torus for each lap down the track (Fig. 2c, Extended Data Fig. 2) (Methods). Consistent with this entrainment, the activity bump took increasingly straighter trajectories as visual landmarks were added (Fig. 2d,e,Extended Data Fig. 4).

**Figure 4.**
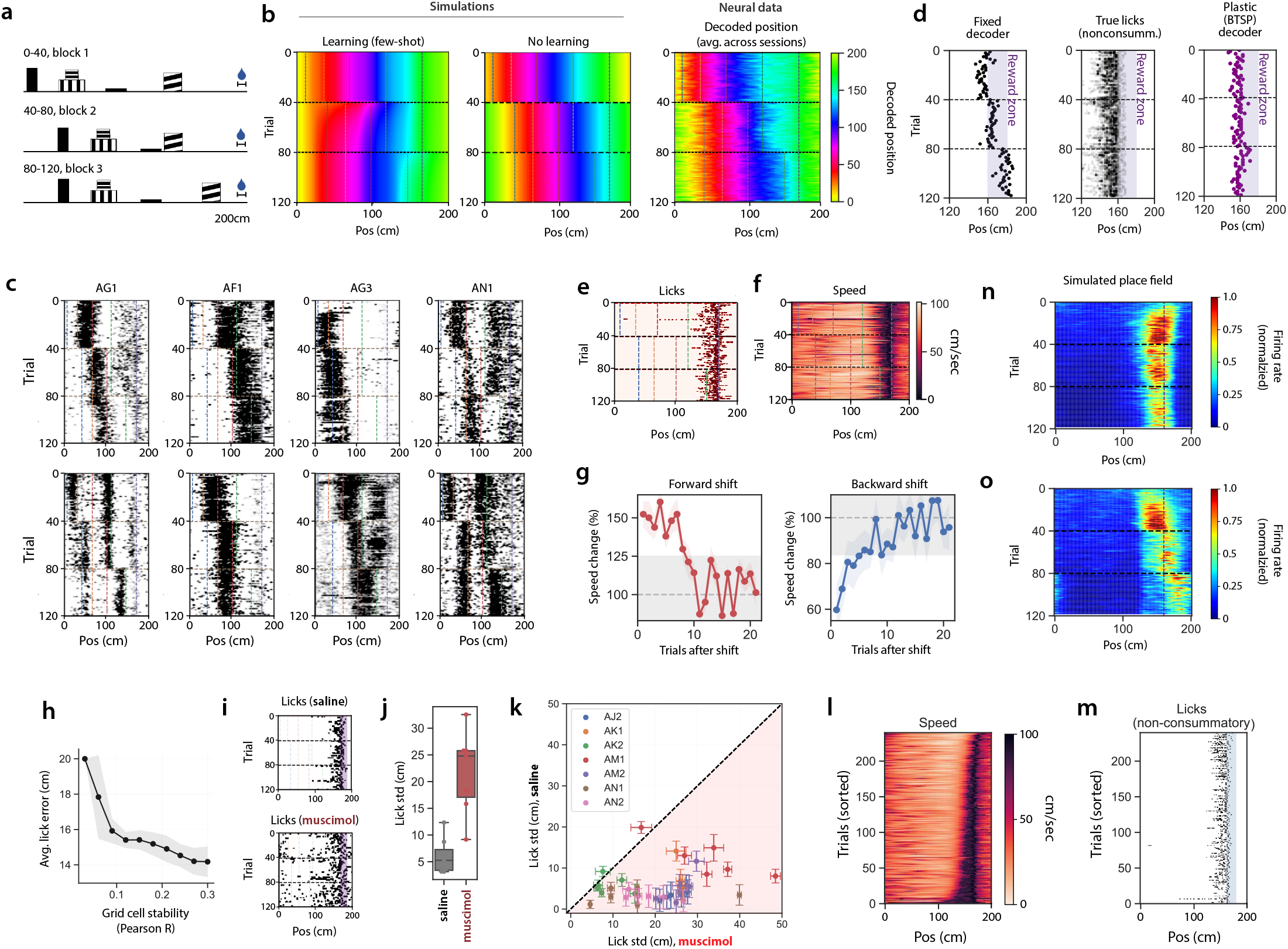
Behavioral time scale plasticity in a downstream circuit could enable rapid spatial learning despite fixed MEC spatial maps. **a**. Schematic of the hidden reward task for the “forward shift condition”. The unmarked reward zone is indicated by the water drop. The mouse traveled a variable distance from the start of each trial to the first landmark. **b**. *Left-Middle*: Decoded position from a population of simulated grid cells which exhibit few-shot learning (*left*) and no learning (*middle*) on the “forward shift condition”. Horizontal black dotted lines correspond to the trial in which the location of the visual cue shifts. Vertical lines indicate the location of the landmarks for each block of trials. *Right*: Decoded spatial position from a linear decoder trained on the population activity of simultaneously recorded grid cells in a tower-shift experiment, averaged over modules, sessions, and animals (animal n = 5, session n = 10, module n = 29, cell n = 1325). **c**. Example grid cell rasters in the hidden reward task from 6 different mice (animal ID on top of each raster). Cues and trial blocks labeled as in (b). **d**. *Left*: Predicted lick locations for a fixed decoder trained on all three blocks. Dots represent licks. *Middle*: Experimentally measured non-consummatory licks across all sessions and mice. *Right*: Predicted lick locations for a plastic decoder trained with a BTSP learning rule. **e**,**f**. Lick (e) and running speed (f) rasters for one recording session. Horizontal and vertical lines as in (b). **g**. Change in running speed in the 10 cm window preceding the reward zone following a single tower shift, averaged across 8 animals and 18 sessions (line indicates mean, colored shaded areas indicate SEM). Behavior quickly recovered to baseline (gray dashed line) after fewer than 10 trials on average. Gray shaded area represents 95% confidence interval of trial-to-trial speed variability in the environment preceding the tower shift. **h**. The average licking error (average distance of non-consummatory licks from reward zone) was large on trials where the grid cell population code was unstable (quantified by average Pearson r correlation to other trials within a block). Licking error decreased on trials where the grid cell code was more stable (shaded area represents SEM across trials, animal n = 8, session n = 17, module n = 50, cell n = 2099). Spatial stability of the grid cell population code, quantified by average trial-by-trial Pearson r correlation of population activity within a block (Pearson r = −0.82, p < 1e-5). **i-k**. Muscimol disrupts task performance. (i) Lick rasters for a session with saline (top) and muscimol (bottom) injection. (j) Licks are significantly less concentrated around the reward zone for mice after muscimol (red) rather than saline (black) injection. Box-and-whisker plot. Horizontal lines represent median and quartiles. Points represent individual animals and sessions (animal n = 7, session n = 7). (k) Across blocks, sessions, and mice, licks are significantly less concentrated around the reward zone for the session after muscimol (red) compared to saline (black) injection. Dots indicate mean, and error bars indicate SEM across 40 trials (animal n = 7, session n = 7). Note, licks are less clustered near the reward location after muscimol injection. **l**,**m**. Licking behavior was not predicted by a timing-based mechanism. (l) Speed raster as in (f) (left), with trials sorted by average speed preceding the reward zone. Speed varies significantly across trials but licking behavior (center) is consistent when plotted in the same sort. (m) Average running speed does not correlate with the licks center of mass (COM). **n**,**o**. Simulated CA1 place cell using a BTSP learning rule on MEC activity from our recordings (Methods). Plots are color coded for minimum (blue) and maximum (red) values. (n) The place cell adapts to distorted grid cell firing patterns to retain a consistent reward-predictive place field across blocks. (o) When plasticity is knocked out, place fields shift the location of their peak firing with the grid cell firing patterns between blocks.

Although grid cells showed less drift with the addition of visual landmarks, the addition of a single landmark was sufficient to induce spatial remapping, such that the shape of the tuning curve of individual grid cells changed (Fig. 2b,f,h). In quantitative terms, the grid cell population maintained high spatial correlation within, but not across, blocks containing at least one VR tower (Fig. 2f) (p < 1e-10, Wilcoxon rank sum test, session n = 14, animal n = 5, module n = 35, cell n = 747). This indicates that grid cells remapped such that they retained little spatial structure from previous trial blocks. Moreover, calculation of a moving window population vector correlation revealed that remapping occurred rapidly between trial blocks (between blocks 6 and 7 it took an average of 1.29 ± 0.07 trials for the grid cell map to more closely resemble the average tuning curve in block 7 than in block 6, Pearson r, session n = 14, animal n = 4, module n = 35, cell n = 747) (Fig. 2f). This number was similar on animals' first exposures to the track (1.42 ± 0.23 trials, session n = 7, animal n = 7, module n = 14, cell n = 328).

To understand the complete remapping of individual grid cells between trial blocks, we again considered grid cells as a population. First, we noted that despite spatial remapping across trial blocks, grid cells maintained low-dimensional continuous attractor dynamics: both temporal correlations (Fig. 2g) (Pearson r, session n = 14, animal n = 4, module n = 35, cell n = 747) and phase relationships (Extended Data Fig. 2) were preserved. When we examined the activity bump’s trajectories, we noted a change in its angle or length across trial blocks (Fig. 2h,i). This provides a remarkably simple understanding of remapping as just taking a new trajectory on a toroidal attractor ^57^, thereby demystifying the complex re-organization of individual grid cells during a remapping event, a process that can appear random aside from the preserved correlations between grid cells.

### One-shot incorporation of landmarks induces small systematic grid distortions

How does the addition of new landmarks to the build-up track (Fig. 2a) rapidly shift the activity bump trajectory to follow a new direction on the neural sheet yet support stable subsequent bump trajectories? One possibility is that landmark inputs rapidly shift where on the neural sheet they project, such that they align with the appropriate location of the bump’s COM on the neural sheet. This alignment would have to be nearly instantaneous between landmark inputs and grid cells. While such learning between landmarks and internal coordinate estimates have been reported in the fly compass system ^18,19^ and, under some conditions, in the mammalian head direction cell system ^58,59^, it can take on the order of minutes to accomplish. Another possibility is that landmark input remains fixed in terms of where it projects onto the neural sheet. This would result in stable but distorted maps. In other words, the distance traveled in physical space may not retain a fixed proportion to the distance traveled by the activity bump on the neural sheet. To consider these two possibilities, we designed an experiment in which mice navigated 12 VR environments, each consisting of the same set of five landmarks (Fig. 3a). For a given block of 20 trials, these five landmarks were placed at random locations, and thus the locations and order of the same landmarks differed across the 12 environments (the “random environment track”, animal n = 14, grid cell n = 4,226) (Fig. 3a).

In these experiments, we performed a series of analyses that considered the bump trajectory on single trials in the presence of visual landmarks. We found that the same forcing function that entrains the bump to periodic trajectories (Fig. 2e) also locally *distorted* the bump trajectory, such that distances traveled in physical space were no longer proportional to distances traveled by the bump on the neural sheet (Fig. 3b-e). This was quantified in two ways. First, we measured the anisometry of the map from the animal’s movements to the movement of the bump on the 2D toroidal neural sheet (Fig. 3b,c). Intuitively, anisometry provides a measure of how stretched or compressed distances on the neural sheet become relative to distances traveled in physical space. We found that the anisometry of the bump trajectory was significantly higher in the presence of landmarks compared to in the dark (Fig. 3c, KS test, p < 1e-10, animal n = 6, session n = 7), indicating the activity bump trajectory was distorted in the presence of visual landmarks. Second, since the animal was constrained to straight-line trajectories, the activity bump should follow a “straight line” on the curved 2D toroidal surface (a geodesic). We found that over short distances (one to a few trials), the bump deviated substantially from geodesic trajectories in the presence of landmarks (Fig. 3d,e). This is analogous to the visual landmarks “tugging” the activity bump off a straight-line trajectory and is consistent with experiments in 2D environments where environmental boundaries locally distort the grid map ^27,31,33,34,36,37^.

While this “tugging” was strong enough to entrain the bump to periodic trajectories (Fig. 2e, Extended Data Fig. 5), it was not strong enough to “pin” the bump to a particular spot on the neural sheet each time the animal passed the corresponding landmark across different blocks (Fig. 3f,g). Indeed, the activity bump’s position was no more concentrated when the animal was near a given landmark versus a random location (Fig. 3g,h). This results from the fact that the bump’s trajectory in a new environment was a function of both the organization of the landmarks and the bump’s history. In other words, the bump could not deviate very far from the trajectory it took on the previous trial. This history dependence was further evidenced in a subset of sessions in which we repeated blocks of trials in a VR environment twice, separated by several blocks of visually distinct environments (build-up track session n = 14, animal n = 4, module n = 35, cell n = 747). We found that grid cells developed different maps in the two repeated environments, despite them being visually identical (∼50% of the time, Extended Data Fig. 4).

### A fixed landmark to grid cell map can predict novel spatial representations

We next asked whether a simple weak landmark pinning model of grid cells provides enough underlying structure to the bump trajectory to *predict* the pattern of distorted grid cell firing activity in a novel environment. To answer this question, we considered the activity of grid cells in 11 of the environments from the random environment track and asked whether we could predict their firing in a 12th held-out environment. We modeled grid cell activity using a simple model of an activity bump moving on a 2D toroidal sheet (Fig. 3f) (Methods). In the absence of landmarks, the model bump took trajectories whose angle drifted according to the diffusion constant D, which we previously extracted from neural data (Fig. 1). When the simulated animal came within 40 cm of a landmark, the model bump was tugged towards the landmark’s “pinning phase” on the 2D toroidal sheet (Fig. 3f). To estimate the pinning phase for each landmark from the neural data, we computed the centroid position of the activity bump on the neural sheet, conditioned on the animal being at that landmark in 11 of the 12 environments (Methods).

After identifying the pinning phase for each landmark, we simulated the bump trajectory on the held-out environment. We used this trajectory to simulate spike trains for each model grid cell by computing a firing rate based on the proximity of the grid cell’s 2D phase to the location of the bump at each time point. The model grid cell spike trains showed a remarkable resemblance to real grid cell spike trains (Fig. 3h-j, Extended Data Fig. 6). To quantify the performance of the model, we measured the average correlation between model grid cell spike trains and neural grid cell spike trains. We found that this simple model was able to predict the firing patterns of many grid cells with a high Pearson correlation across held-out environments, and that the best-predicted cells were also the most stable (Fig. 3i). We then compared these correlations to two shuffled conditions: randomly shifting the simulated tuning curves (Fig. 3h), and scrambling the landmark locations used to train the model (Extended Data Fig. 6). The 2D dynamical model consistently outperformed both null models.

Motivated by prior theoretical work and experimental evidence from *Drosophila* ^18–21,60^, we incorporated a Hebbian learning mechanism with a tunable learning rate into our model. Consistent with our previous results, we found the learning rate that best explained the neural data was zero, suggesting that grid cell firing activity across novel environments was best explained by a fixed, non-plastic circuit (Extended Data Fig. 6).

### One-shot MEC maps enable flexible adaptive behavior despite small distortions

A fixed landmark to grid cell network allows for rapid, stable maps to form even when new landmarks are observed, which could support behaviors that occur on short timescales. However, such fixed networks also induce distortions to the regularity of the grid map. This raises the question: how well can animals navigate with a distorted grid map? To investigate this question, we developed a VR task in which animals had to use a combination of distance run and landmark positions to find a hidden reward zone (the “hidden reward task”).

Animals were trained on a 200 cm long track (Fig. 4a), consisting of four visual landmarks and an unmarked 10 cm wide reward zone that started 20 cm past the last tower (baseline block). Animals could lick anywhere within the reward zone to trigger a water reward. To maintain running behavior, animals received an automatic reward at the end of the reward zone if they failed to lick within the reward zone. To prevent the animals from using time elapsed or distance run since the last reward, we varied the distance from the beginning of the track to the first tower randomly on each trial. We found that after training, animals licked prior to the reward zone and slowed their running speed after the last tower, indicating they had learned the location of the hidden reward zone (Extended Data Fig. 7).

After training, we recorded neural activity during the hidden reward task on the same linear track. In the first block of 40 trials, the four landmarks remained in the same location as in the training sessions. In the second block of 40 trials, we moved the first three landmarks forwards (towards the rewarded location) by 30 cm. In the final block of 40 trials, the last landmark was shifted forwards by 30 cm. We refer to this as the “forward shift condition”. To ensure that the behavioral effects we observed were symmetric, we repeated these three blocks in reverse, shifting a subset of landmarks backwards (towards the beginning of the track) on each subsequent block. We refer to this manipulation as the “backward shift condition” (Fig. 4a, Extended Data Fig. 7).

Based on our previous observations, such tower shifts should induce a distortion in the grid map. However, if the animal estimates its spatial position from the grid map, then the grid map must recover from any landmark induced distortion, or the animal will lick too early/late when the towers are shifted forward/backward and miss the reward zone. We simulated the possible outcomes of this experiment with and without learning, using the same model as in Figure 3. With learning (Fig. 4b, left) the simulated animal’s decoded position was deflected by the landmark shift, but recovered after a few trials (i.e. few-shot learning condition). Without learning (Fig. 4b, middle), the simulated animal’s decoded position was permanently deflected and never recovered. We then repeated this procedure for our experimental data, by training a linear decoder on the population activity of grid cells recorded during the hidden reward experiment. Consistent with the results from the random environment track the neural data (Fig. 4b, right) more closely resembled the simulated no-learning outcome (Fig. 4b, middle): the animal’s estimated position based on the decoded activity of the grid cell population deflected significantly with the landmark shift and never recovered. This deflection could also be clearly observed in single-grid cell activity across animals and sessions (Fig. 4c). Furthermore, we found that *no fixed linear decoder* could correctly decode the animal’s position across blocks, including a decoder trained with neural activity from all three blocks (Fig. 4b, right). The resulting decoder predicted that the animal would lick in very different spots before and after the landmark shift (Fig. 4d). Remarkably, however, the animals rapidly learned to adapt their behavior to the deformed environment. Unlike the neural activity, behavioral performance was only momentarily deflected and animals returned to their previous performance accuracy after fewer than 10 trials in the new environment (Fig. 4e-g). This was the case for both forward and backward shift conditions.

What could explain this difference between the grid map and the animal’s behavior? One possibility is that grid cells are simply not involved in this path integration task. However, three pieces of evidence support a role for grid cells in this task. First, grid cell spatial stability was strongly correlated with behavioral performance (Fig. 4h). Second, bilateral muscimol injection into MEC significantly silenced neural activity (Extended Data Fig. 1, 8) and impaired behavioral performance compared to a comparable saline injection (Fig. 4i-k, Extended Data Fig. 8). In a recording where muscimol did not appear to fully silence MEC, we found that behavioral performance was also not fully impaired (Extended Data Fig. 8). Third, distance-based strategies were more predictive of the animals’ behavior than time-based strategies (Fig. 4l,m). If animals used a time-based mechanism to locate the hidden reward, then they should lick late on trials where they run quickly and vice versa. However, we did not observe this type of relationship, consistent with the hypothesized role of grid cells in supporting distance-based navigation strategies ^25^.

Another possibility is that while landmark connectivity to grid cells is fixed, grid cell connectivity to a downstream decoder supporting behavior might be plastic. To explain behavior, such a decoder would have to adapt to an altered grid cell code after just one or a few trials. Recent experimental work has identified a rapid, behavioral timescale synaptic plasticity (BTSP) mechanism that governs place field formation in CA1 and is driven by a signal from MEC ^8,61,62^. To test the feasibility of this mechanism, we trained a linear decoder using the BTSP learning rule on grid cell activity from our recordings (Methods). Unlike the fixed decoder, this plastic decoder was able to quickly adapt to landmark shifts, and could reliably predict the reward location across all environments (Fig. 4d, right). Moreover, the plastic decoder makes an explicit prediction for how a hippocampal place cell’s firing field would respond to the tower shift experiment if it adapted according to a BTSP learning rule (Fig. 4n), and how place cells would respond if plasticity between MEC and CA1 were knocked out (Fig. 4o).

## Discussion

We used Neuropixels recordings of thousands of grid cells in mice navigating diverse VR environments to show that MEC is capable of building *stable*, consistent maps of novel environments after just a single exposure (one-shot). Spectral analyses revealed that this was accomplished by visual landmarks driving a bump of neural activity onto stable periodic trajectories on a 2D toroidal neutral sheet with attractor dynamics. However, visual landmarks also *distorted* the grid map, such that distances traveled by the animal in real space were no longer proportional to distances traveled by neural activity on the neural sheet. Nevertheless, these distortions followed quantifiable dynamics, which allowed us to use a simple 2D dynamical model to *predict* grid cell responses to novel arrangements of visual landmarks. Together, these results point to grid cells as providing a map which, though distorted, is rapid and stable enough to subserve one- or few-shot learning in novel environments via a downstream decoder that can implement a biologically plausible learning rule ^8,62^ .

These findings are consistent with prior works demonstrating attractor dynamics in the MEC ^41–49,57^ and observations of grid map distortions in 2D and 3D environments ^30–33,36,37,63^. Our results here suggest that such grid map distortions are the result of fixed, non-plastic and weak grid attractor pinning from landmark inputs, providing a unified framework for interpreting the repertoire of MEC grid cell remapping observations in response to changes in environmental features. This fixed and weak landmark to grid map pinning system represents a solution between the extremes of a purely idiothetic based system, in which landmarks have no influence, and one in which landmarks strongly pin the grid cell system in the absence of plasticity. In the latter case, the distortions caused by strong landmark pinning would likely fall in a category too large for correction even by a plasticity rule such as BTSP, and therefore would also require plasticity from landmark cells to grid cells to smooth out large distortions from strong pinning ^21^. Such requisite plasticity would preclude one-shot learning of novel environments capable of subserving flexible behavior, which is a key computational advantage of the weak landmark to grid cell pinning solution. Indeed in this one-shot environmental mapping solution, our data show that the influence of novel landmark features on the grid map causes some distortion, but these distortions remain within the operating time regime of the BTSP model, allowing rapid incorporation of novel landmark features without major map reorganization.

Our observations of fixed and weak pinning from landmark inputs differ from work in *Drosophila* showing learning between landmark inputs and the fly compass system, which exhibits homologous dynamics to mammalian attractor network based navigation systems ^18–20,43,45,48,49,53,58,64,65^. In *Drosophila*, Hebbian-like plasticity is hypothesized to underlie the association between visual neurons and compass neurons that are co-active, thus allowing realignment of the fly compass system on the order of minutes when visual landmarks change ^18,19,66^. However, there are several possibilities for the differences in the fly orientation and mammalian grid cell system. First, *Drosophila* and rodents face different ethological pressures. Fruit flies do not have a permanent home location, while mice often explore novel environments in addition to having a more permanent home location. This could require mice to retain the ability to rapidly create reliable maps of any novel environment such that they can take a direct trajectory back home ^40^. Consistent with this ethological hypothesis, invertebrates that do have home locations (such as ants and honeybees) struggle to navigate to goal locations after alterations to spatial landmarks ^67,68^. Second, the plasticity mechanism in the *Drosophila* compass system may differ from the mammalian grid cell system. Plasticity in the *Drosophila* compass system results in weakening of the (inhibitory) synapses between associated visual landmark inputs and compass neurons. This may differ from the plasticity rules thought to operate in the mammalian navigation system, with associations between visual landmark inputs and grid cells strengthening rather than weakening ^20,21,29^. Moreover, recent work has revealed that the learning rate of plasticity in the *Drosophila* compass system depends on dopamine ^69^, but whether such a mechanism exists in MEC remains completely unknown. Third, a closer parallel could exist between *Drosophila* and the mouse when considering not one, but multiple, networks. The mammalian system exhibiting the closest homologies to the *Drosophila* compass system may be earlier in the hierarchy of navigational coding, for example, the head direction system in the antero-dorsal thalamus ^58,65,70,71^ or the dorsal tegmental nucleus, the earliest known processing of head direction in the mammalian brain ^72^. Another possibility is that the expansion in the number of neurons in the mammalian cortex enabled the formation of dedicated circuits with differential preferences for stability or flexibility. Thus, the mammalian navigation system may accomplish the same computational algorithms as the *Drosophila* system but with the MEC prioritized for stable, rapid, inflexible maps with small distortions and the hippocampus prioritized for flexible processing of novel landmark features.

The specific nature of landmark inputs to the grid cell neural sheet remains an open question. Theoretically, this input could be provided by any neuron that reliably encodes information about a landmark, for example, non-grid spatial ^73,74^, border ^2,6^ and object vector cells ^5^ in MEC, neurons in V1 that fire near visual landmark features such as towers in VR environments ^25,75,76^, or boundary vector ^77^ and corner coding cells ^78^ in the subiculum. The model we utilize here assumes a simple connectivity structure between landmarks and grid cells, where landmarks have a preferred “pinning phase” on the neural sheet. However, future work could consider whether more general and unstructured connectivity patterns between landmark-tuned cells and the grid cell sheet, which would pin not to a single “phase” but to a “field”, could accomplish the same effect of driving the bump of activity onto stable trajectories.

While the current results point to the MEC grid map as relatively inflexible over short behavioral timescales, our results do not rule out the possibility that Hebbian plasticity from sensory inputs to grid cells can alter the grid map over longer time scales. Consistent with this idea, grid cells form global maps for connected environments on the order of many days ^28^. Across days, grid maps tend to exhibit stable spatial firing patterns for the same environment ^1^, suggesting that landmark features may set the initial phase of the activity bump on the neural sheet when an animal enters a familiar environment. One possibility is that plasticity between landmarks and the grid cell map is computationally and metabolically expensive and thus only occurs when it is adaptive. This could include, for example, scenarios in which environments are highly familiar, such as a home base, or could occur for landmarks with high valence or task relevance, which may impose stronger pinning dynamics on the grid cell neural sheet ^32,33,79,80^.

In summary, accurate internal estimates of location must be derived by fusing information from the integration of idiothetic self-motion with information derived from external landmarks. The fusion of these two distinct information sources becomes even more computationally challenging in dynamic environments where putative landmarks can themselves move over behaviorally relevant time scales. In the design space of algorithms for fusing self-motion with landmarks, our findings reveal that the combined mammalian grid cell-landmark system achieves the best of both worlds between two extremes via a weak, fixed landmark to grid cell pinning solution. One extreme of no landmark pinning obviates the need to deal with the complex problem of dynamic landmarks, but precludes the creation of stable spatial maps under the noisy integration of self-motion alone. The other extreme of strong landmark pinning would require plasticity between landmarks and grid cells to smooth out substantial grid cell distortions, thereby precluding the possibility of the rapid one-shot creation of such stable maps with small distortions that can instantly map landmark rearrangements or completely novel environments in a single exposure. The weak, fixed, landmark to grid cell pinning solution favored by MEC is thus a computationally powerful approach enabling one-shot learning of grid maps in new or dynamic environments, with small distortions, that can then subserve flexible adaptive behavior through downstream plasticity.

## Methods

### Experimental model and subject details

All techniques were approved by the Institutional Animal Care and Use Committee at Stanford University School of Medicine. A total of 19 female C57BL/6 mice were used for this paper. Electrophysiological recordings were conducted on 15 female C57BL/6 mice aged 12-24 weeks at the time of the first surgery (headbar implantation). Mice were housed with littermates on a 12-hour light-dark cycle. After the craniotomy surgery, mice were housed singly to prevent disturbance of the surgical site. An additional 4 female C57BL/6 mice were used for fluorescent muscimol histology only.

### Virtual reality (VR) environments

The VR recordings setup was nearly identical to that in ^25^. Mice were head-fixed and ran on a 15.2 cm diameter foam roller (ethylene vinyl acetate) constrained to rotate about one axis. Rotation of the cylinder was tracked with a rotary encoder (Yumo 1024 P/R) and relayed to a microcontroller (Arduino UNO) using custom software. The virtual reality environment was developed using custom code in Unity 3D. The virtual scene was displayed on three 24-inch monitors surrounding the mouse. Water rewards were delivered through Tygon tubing attached to a custom-built lickport. The lickport consisted of a lick spout and an infrared (IR) light beam to detect licks. Water was delivered via opening of a solenoid valve. Phase 3B Neuropixels 1.0 silicon probes ^51^ were mounted on a custom-built rotatable mount and positioned using a micromanipulator (Sensapex uMp). Virtual tracks specific to each task are detailed below.

#### Build-up track

The build-up track was 320 cm in length and upon reaching the end of a track, mice were seamlessly teleported to the beginning of the track without any visual discontinuity. The experiment consisted of nine blocks. The first and last blocks occurred in the dark for a minimum of 10 minutes each, where no water rewards were given and no visual features were displayed. Blocks two through seven consisted of 40 trials each, and block eight consisted of 20 trials. In the second block, water rewards were automatically dispensed at the end of the track. In the third block, a repeating checkerboard optic flow pattern was displayed on the ground. In the fourth through seventh blocks, visually distinct tower landmarks were cumulatively added. In the eighth block, the tower landmark added in the previous block (block 7) was removed. All tower landmarks were displayed on either side of the animal. The visual lookahead was 300 cm.

#### Random environment track

The random environment track was 100 cm in length with a 100 cm visual lookahead. The experiment consisted of 14 blocks. The first and last blocks occurred in the dark and consisted of at least 100 trials where no water rewards were given and no visual features were displayed. In the second through thirteenth block, consisting of 20 trials each, a set of five familiar visual landmarks were pseudo-randomly rearranged. All visual landmarks were visually distinct from one another. Three of the landmarks were tower landmarks, and two were ground landmarks. One of the ground landmarks was associated with a water reward. For each random rearrangement, the distance between any two landmarks had to be at least 15 cm apart and the position of the first landmark had to be at least 15 cm from the start of the track.

#### Hidden reward track

The hidden reward track was 200 cm in length with a 100 cm lookahead. The track consisted of the same visual cues in the random environment track except for the absence of the reward cue. Thus, there were four visual cues on the track (three tower landmarks and one ground landmark). The positions (in cm) of the landmarks were 40, 65, 100, and 145 in the baseline condition. A 10 cm hidden reward zone was positioned at 170 cm from the start of the track. Animals could lick anywhere between 160 cm −170 cm to trigger a water reward. If an animal failed to lick within the zone, they received an automatic reward at 170 cm. To encourage animals to pay attention to their position relative to landmarks, and not simply path integrate a fixed distance between rewards, an offset value was drawn from a uniform distribution between 0 cm and 30 cm on each trial. This offset value was added to the positions of each of the landmarks and the reward zone, resulting in a variable inter-trial distance before encountering the first landmark. This offset value is not shown in Figure 4 for visualization clarity. The experiment consisted of seven blocks: A, B, C, D, C, B, A. Block A was the same landmark arrangement as the training track and served as the baseline condition. Block B was identical to A, except the last visual landmark was moved 35 cm closer to the start of the track. Block C was identical to block B except the first three landmarks were also moved 35 cm closer to the start of the track. Block D is a dark block where no water rewards were dispensed and no visual cues were displayed. For the second set of shifts (C, B, A), the landmark arrangement in Block C was used as the baseline condition. All blocks except block D consisted of 40 trials each. Block D consisted of at least 100 trials.

### Behavioral training and handling

Prior to headbar implantation, mice were given an in-cage running wheel (Bio-Serv K3570 and K3251). To ensure ease of use, the running wheel pin was manually shaved to reduce friction. Mice were handled for at least a day before headbar implantation. Handling included holding each mouse for at least half a minute before setting them on the running wheel to encourage their use of it.

After headbar implantation, mice were injected with Rimadyl (5 mg/kg) and Baytril (10 mg/kg) daily for three days. Then they were restricted to 0.8 mL of water per day. Mice were weighed daily to ensure they stayed above 80% of their baseline weight. After at least one day of water deprivation, mice began training on the virtual reality setup. All mice received the same initial basic training. The first stage of training acclimated mice to head-fixation and running on the VR wheel. No VR visual cues were presented on the monitors. The VR wheel was manually moved to encourage walking on the setup. As the wheel was moved, a 1 mL syringe with water was manually and simultaneously brought closer to the mouse. After a half revolution of the wheel, the syringe dispensed water.

After mice could run on their own (> 1/2 revolution of the VR wheel), the second stage of training began. This stage acclimated mice to a lickport, which dispensed water (∼1.5-2 μl at a time) and detected licks with an infrared (IR) beam sensor. The dispensation of water was associated with the sound of a solenoid valve opening. After mice reliably licked upon hearing the solenoid click, they proceeded to task-specific training, as described below.

#### Build-up track

mice continued training in the absence of any virtual reality visual cues. They were trained on a training track that was completely dark. The track contained an automatic water reward that was given after a certain distance was run. This distance was increased adaptively within and across training sessions to encourage consistent running in the dark. To avoid training mice to encode any particular distance, the average distance required to receive a water reward was different from session to session, ranging between 400 cm to 1000 cm between rewards. Mice were considered ready for recordings when their average speed exceeded 250 virtual cm / s for several days.

#### Random environment track

mice were introduced to the five visual cues that would be used in the experiment by running on a VR track with a specific arrangement of the cues for 100 trials per day for three consecutive days. Then, the day before recording, mice were first trained with the same arrangement of cues for 20 trials followed by 11 blocks of 20 trials each where the cue locations were pseudo-randomly shuffled. This shuffling of cues was done to ensure mice still licked at the reward cue. If a mouse did not complete 100 trials in less than an hour, the original training was completed with the same initial arrangement of cues. Mice were ready for recordings when they could run 100 trials in less than an hour.

#### Hidden reward track

upon completion of recordings in the random rearrangement experiment, 7 mice were used for the hidden reward experiment. The visual cues in both tracks were identical except for the absence of a visible reward cue in the hidden reward experiment. Training proceeded by giving mice an automatic water reward at the end of a hidden reward zone. Initially, the hidden reward zone was 30 cm in width so that any licks within 30 cm of the end of the zone would trigger a water reward to be dispensed. The reward zone was gradually decreased in size over training sessions until it was 10 cm in width. Mice were ready for recordings when they triggered water rewards with licks within the reward zone prior to the automatic reward on at least 70% of trials.

### *In vivo* survival surgeries

For all *in vivo* survival surgeries, a mixture of oxygen and isoflurane was used for induction (4%) and maintenance (0.5%-1.5%) of anesthesia. After induction, buprenorphine (0.05-0.1 mg/kg) was injected subcutaneously. Upon completion of surgery and for three additional days after, Baytril (10 mg/kg) and Rimadyl (5 mg/kg) were subcutaneously injected.

#### Headbar surgery

The first surgery consisted of attaching a stainless steel headbar, implanting a gold ground pin, and making fiducial marks on the skull. The headbar was cemented to the skull with Metabond (Parkell S380). The ground pin was implanted roughly 2.5 mm anterior and 1.5 mm lateral of bregma. Fiducial marks were made at +/-3.3 mm relative to midline and 3.7 mm posterior of bregma. These marks served to guide insertion of Neuropixels probes and microsyringes for muscimol infusion.

#### Bilateral craniotomies

Following completion of training, bilateral craniotomy surgeries were conducted. Craniotomies removed a small portion (∼200 μm along the medial-lateral axis and ∼300 μm along the anterior-posterior axis) of skull posterior to the fiducial marks and exposed the transverse sinus. Craniotomies were covered with a silicone elastomer (World Precision Instruments, KWIK-SIL). Then a small plastic well was implanted over each craniotomy and affixed with dental cement.

#### Microsyringe infusions

For the hidden reward task, bilateral injections of muscimol (Sigma-Aldrich M15223) or saline control in MEC were performed in seven mice weighing 18-25 g. Additionally, two of the seven mice also received BODIPY TMR-X fluorescent muscimol (ThermoFisher M23400) injections on a separate day to confirm the neural and behavioral effects of fluorescent muscimol compared to untagged muscimol. An additional four mice received bilateral injections of fluorescent muscimol in MEC without behavior or Neuropixels recording to confirm appropriate microinfusion targeting. See Extended Data Table 1 for a detailed list of all the mice in this study, which sessions they ran, and which drugs they received.

1 mg of muscimol or 1 mg of BODIPY TMR-X muscimol conjugate was diluted in 4.3 mL and 0.82 mL, respectively, of filtered 1X phosphate-buffered saline (PBS) (Fisher Scientific BP3991), resulting in a final concentration of 2.0 mM.

Prior to infusion of muscimol or saline control, craniotomies were inspected and cleaned with sterile saline if needed. Bilateral infusions were achieved using a 35-gauge needle (World Precision Instruments NF35BV-2) and a 10 μL Hamilton Syringe (World Precision Instrument NANOFIL). The syringe was angled 14 degrees from vertical. Then, the tip of the needle was positioned +/-3.3 mm medial-lateral (ML) relative to the midline, 100-150 μm in front of the transverse sinus, and touching the dura surface. Then, the syringe was advanced into the brain at a rate of 8.3 μm/s using a Neurostar robot stereotaxic. 220 pmol or 110 nL of either form of muscimol or saline control was injected at 334 nL/minute using an ultra-micropump (World Precision Instruments UMP3) at each of four sites along the dorsoventral axis the MEC. These sites were 1200 μm, 1950 μm, 2700 μm, and 3450 μm below dura. Following injection at each site, the syringe was left in place for 1 minute to allow for sufficient diffusion of the drug.

### *In vivo* electrophysiological data collection

All recordings took place after the mice had recovered from bilateral craniotomy surgeries (a minimum of 16 hours later). Immediately prior to recording, the Neuropixels probe was dipped in one of three dyes (DiD, DiI, DiO, ThermoFisher V22889). The probe was dipped 10-15 times with 10 seconds in between dips. If three recordings had already been performed in one hemisphere of the brain, no dye was used for subsequent recordings in that hemisphere. Then, mice were head-fixed on the VR setup, and the craniotomy sites were exposed and rinsed with saline. The Neuropixels probe was angled 12-14 degrees from vertical and positioned using the fiducial marks made during the headbar surgery and 50-200 μm anterior of the transverse sinus. The medial-lateral position of the probe relative to the medial edge of the fiducial mark, the anterior-posterior position relative to the transverse sinus, and the angle relative to vertical were all recorded for replicability and probe localization (see “Histology and probe localization” below). The 384 active recording sites on the probe were the ones occupying the 4 mm closest to the tip of the probe. The reference and ground were shorted together and the reference electrode was connected to the gold pin implanted in the skull. Then, the probe was advanced into the brain using a Sensapex micromanipulator, typically at 5 μm/s but no faster than 10 μm/s. Insertion of the probe continued until resistance was met or until several channels near the tip of the probe were quiet. Then, the probe was retracted 100-150 μm and allowed to settle for at least 15 minutes prior to recording. Finally, the craniotomy site was covered in silicon oil (Sigma-Aldrich 378429).

Raw signals were filtered, amplified, multiplexed, and digitized on-probe. SpikeGLX (https://billkarsh.github.io/SpikeGLX/) was used to sample voltage traces at 30 kHz and filter between 300 Hz and 10 kHz for the AP band. SpikeGLX was also used to acquire auxiliary signals. Each Unity VR frame emitted a TTL pulse, which was first relayed to an Arduino UNO, which then relayed it to an auxiliary National Instruments data acquisition card (NI PXIe-6341). These auxiliary signals were used to synchronize VR data with electrophysiological traces.

At the end of each recording, both craniotomies were inspected, rinsed with saline, and then covered in Kwik-Sil. Animals were then placed back in their home cage and given additional water to raise their body weight at least 1 gram higher than their weight at the start of the day. This helped ensure their body weight never dropped below 80% of their starting weight prior to water restriction. The probe was rinsed and soaked for at least 10 minutes in a 2% solution of Tergazyme (Fisher Scientific 16-000-116) in deionized water and then rinsed with deionized water alone.

### Histology and probe localization

After the last recording session, mice were euthanized with an overdose of pentobarbital and transcardially perfused with PBS followed by 4% paraformaldehyde (PFA). Then, brains were extracted and stored in 4% PFA for at least 24 hours and then transferred to 30% sucrose solution for at least 24 hours. Finally, brains were frozen and cut into 100 μm sagittal sections with a cryostat, stained with DAPI, and imaged using widefield microscopy (Zeiss Axio Imager 2). All sections that contained dye fluorescence were aligned to the Allen Mouse Brain Atlas using software written in MATLAB ^81^. Then, probe tracks were reconstructed by localizing dye fluorescence in each section. Probe locations for recordings without dyes were estimated relative to previous recordings, using two common reference locations: medial-lateral distance relative to the medial edge of the fiducial mark, and distance anterior of the anterior edge of the transverse sinus.

### Offline spike sorting

Spike sorting was performed offline using the open source spike sorting algorithm Kilosort2, which is a modified version of the original Kilosort algorithm ^82^. Clusters were manually inspected and curated in Phy (https://github.com/cortex-lab/phy). Clusters containing fewer than 500 spikes were discarded. Clusters were also discarded if their peak-to-peak amplitude over noise ratio was less than 3, where noise was estimated by calculating the standard deviation of a 10 ms window preceding each spike. All remaining clusters were manually examined.

VR data was then synchronized to spiking data using the TTL pulse times from Unity and the recorded TTL pulses in SpikeGLX. Custom code written in MATLAB was used to synchronize the data. Confirmation of appropriate syncing was accomplished by noting a high correlation between VR time differences and the corresponding TTL time differences recorded in SpikeGLX. Finally, because the VR frame rate was not constant, we used interpolation to resample behavioral data at a constant 50 Hz, similar to ^25^.

### Identification of grid cells

We identified putative grid cells by computing spectrograms for each neuron’s spike train in the dark, binned in 2 cm spatial bins, using a window of size 3 −5 m. We then extracted co-modular grid cells, neurons that shared the same grid spacing and orientation but differed in phase, by applying k-means clustering to the spectrograms (using k = 5 or 6). We identified a cluster as a putative grid cell module if the average spectrogram correlation (Pearson r) was greater than 0.3 across windows in the first block of trials (n = 15,342 out of 68,484 neurons). We found that these correlations were preserved over much longer timescales, from the first block of dark trials at the beginning of the session to the final block of dark trials at the end of the session (Extended Data Fig. 2).

### Sorting grid cells onto a neural sheet

To sort the grid cells within a module onto a functional neural sheet, we used a standard peak-finding algorithm (scipy.signal.find_peaks) to identify the largest three peaks in the spectrogram within each window, *f*_1_ · *f*_2_, *f*_3_. For each neuron, we then extracted the phases 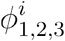 of the Fourier transform at the location of each peak:

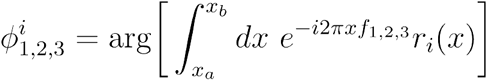

where *r*_*i*_(*x*) is the firing rate of neuron and represents the total distance traveled in the dark. *x*_*a*_,*x*_*b*_ represent the start and end of a window, with the same size as above. We selected the two phases which were best correlated across windows in the first block of dark trials, combined them into a set of N 2D vectors 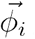 and used them to sort the grid cells onto a functional sheet. To visualize population activity at an instant in time in this sort (Fig. 1e,f), we quantized the phases onto a 2D grid, and plotted activity as a heatmap on this grid. Because this analysis requires good coverage of the 2D phase space, we restricted this analysis to sessions where 36 or more grid cells were recorded simultaneously within a module.

To extract the moment-by-moment attractor state 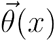 of the grid cell population activity we estimated the center of mass of the bump of activity on the sheet, which can be found by first computing,

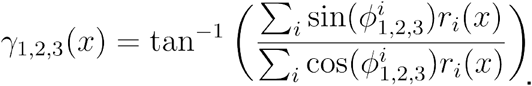

If the population activity of grid cells is well described by the motions of a bump on a 2D sheet, then the three coordinates *γ* _1,2,3_ (*x*) represent only two degrees of freedom. In Extended Data Fig. 2 we show that *γ* _1,2,3_ (*x*) does not take on all possible values, but is instead restricted to a 2D subspace. We can therefore extract these two degrees of freedom, *θ*_1_, *θ*_2_, which represent the center of mass of the bump of activity on the sheet,

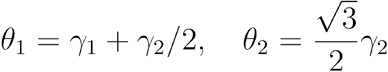

We can visualize the trajectory of the bump of activity on the sheet by plotting the two coordinates *θ*_1_, *θ*_2_ as the animal traverses the VR environment. We plot *θ*_1_, *θ*_2_ for one full session as a heat map over spatial bins and trials in Fig. 2c. Since *θ*_1_, *θ*_2_ are periodic quantities on the torus, we also occasionally unwrap them to plot them as continuous trajectories, plotting them as “revolutions in *θ*_1_ and *θ*_2_” (Figs. 1i, 2d,h).

This procedure for extracting the moment-by-moment attractor state has an equivalent geometric interpretation, as projecting the N-dimensional neural activity onto a 6-dimensional subspace spanned by the vectors *u*_1,2,3_ =sin(*ϕ*_1,2,3_) and *υ*_1,2,3_ =cos(*ϕ*_1,2,3_) (Fig. 1g). In this subspace the neural activity lies close to a manifold with the topology of a twisted torus, and when projected onto each pair of axes, the neural activity traces out a ring (Fig. 1j). Extracting the position *γ* _1,2,3_ (*x*) of activity on any one ring can be performed by computing:

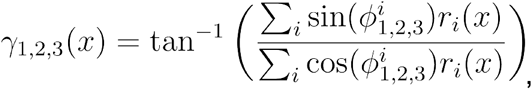

which is exactly the same calculation we used to obtain the 2D attractor state above. Hence extracting the moment-by-moment attractor state is equivalent to identifying the position of the neural activity vector on this torus.

### Remapping between VR blocks

To quantify remapping between blocks in the build-up track environment, we computed the spatial correlation of grid cell population activity 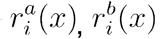, for all pairs of trials *a*,*b*. Averaged across modules, sessions, animals, the resulting correlation matrix (Fig. 2f, bottom) revealed that population activity was stable within a block, but differed between consecutive blocks. The sharp boundaries indicated that remapping happened quickly. To quantify the rate of remapping, we additionally computed the average correlation within a sliding window of 5 trials (Fig. 2f, top),

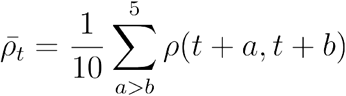

where *ρ*(*a*,*b*) represents the spatial correlation between trials *a* and *b*. Finally, to visually inspect the differences in tuning curves between consecutive environments at a single-cell level, and within a single session, we compared single-cell spatial tuning curves averaged over all 40 trials within the Tower 2 environment (Block 6) and the 40 trials within the Tower 3 environment (Block 7) (Fig. 2h).

### Distortions to the grid cell map in the presence of landmarks

We used two quantifications to measure distortions to the grid cell map in the presence of visual landmarks: anisometry (Fig. 3c) and geodesic curvature (Fig. 3d). A map *θ* is isometric if it preserves distances, i.e. |*x* − *y*| = *d*_map_ (*θ*(*x*), *θ* (*y*)). We defined distance on the grid cell map as distance on the neural sheet 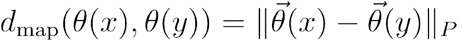, where the subscript *P* indicates the periodic toroidal boundary conditions, accounted for as in ^83^. For each 2 cm interval the animal traveled on the VR track, Δ*x*, we computed the distance the bump of activity traveled on the neural sheet *d*(*x*) ≡ *d*_map_ (*θ*(*x*), *θ* (*x* + Δ*x*)). We then measured the variation in this quantity. Since *d*(*x*) will differ between different modules by an overall gain on average, we quantified variation in *d*(*x*) by the coefficient of variation, over windows of length 16 meters. We found that the coefficient of variation was significantly higher in VR environments than in the dark, indicating that distances on the neural map are distorted in VR.

The second quantity we use to measure distortions to the grid cell map in the presence of visual landmarks is geodesic curvature (Fig. 3d). Since the animal travels in a straight line on the treadmill through the VR environment, if the grid cell map faithfully captures the animal’s trajectory, the bump of activity should travel in a straight line trajectory on the neural sheet, or, equivalently, the neural activity should trace out a geodesic trajectory on the torus (a geodesic being the generalization of straight line on a curved manifold). Parameterizing the 2D torus by the coordinates *θ*_1_, *θ*_2_ the geodesic curvature *κ*_*g*_ reduces to the planar curvature:

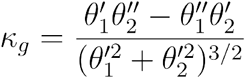

To handle periodic boundary conditions we first unwrap the coordinates *θ*_1_, *θ*_2_ before computing *κ*_*g*_. To extract a single scalar *D* measuring the curvature of a single-trial trajectory, we integrate the geodesic curvature along the trajectory as the animal completes one lap down the VR track:

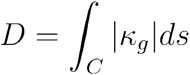

We compute the single-trial geodesic curvatures for trials in the dark and trials in VR, and find that trials in VR have significantly higher geodesic curvature (Fig. 3d).

#### Pinning strength

To quantify the local effect of a VR landmark on the bump trajectory in the random environments experiment, we computed the trial-by-trial position of the bump of activity, conditioned on the animal being at the location of one of the VR landmarks 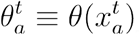, where 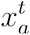 represents the position of landmark on the VR track *a* on trial *t*. 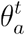 is plotted across trials in one session in Fig. 3g for all five VR landmarks. We then considered the extent to which landmarks were “strongly pinning”, in the sense that the bump of activity was pinned to the same location on the neural sheet each time the animal passed a given landmark (Fig. 3h). To quantify this, for each landmark we measured the dispersion of 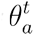 around its mean 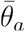 across trials:

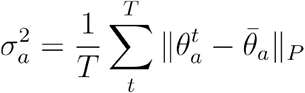

We found that no landmark location was “strongly pinning” compared to a null model conditioned on random locations (gray histogram, Fig. 3h).

### 2D dynamical model

We model the activity of a population of grid cells by a bump moving on a periodic sheet, with the topology of a twisted torus. In the absence of landmarks, the bump moves in straight lines with a velocity 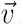. We simulate the effect of visual landmark *i* on the grid cell code as “pinning” the bump of activity to a preferred location 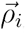,

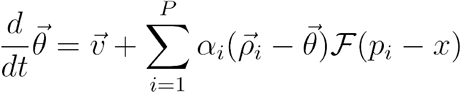

where *x* is the position of the animal on the track, *p*_*i*_ is the location of landmark *i* on the VR track, 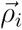 is the preferred 2D pinning location of that landmark on the grid cell sheet, and *α*_*i*_ represents the landmark’s pinning strength. ℱ parameterizes the range of influence of landmarks on the grid cell code. We found that the precise functional form of ℱ did not make a crucial difference, so we chose a particularly simple form: ℱ(*p*_*i*_ − *x*) = Θ (25 − |*p*_*i*_ − 25 − *x*|), where Θ is the Heaviside function, so that each landmark only influenced the grid cell code in the 50 cm leading up to its location on the VR track *p*_*i*_. and *α*_*i*_ the strength of each landmark’s pinning. Finally, we model the velocity as a straight-line trajectory with angle, 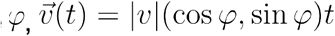, where the slice angle drifts with time at a rate *δ*. The magnitude (or gain) |*υ* | governs the grid scale.

We estimated |*υ* | and *δ* using the responses of the grid cells in the dark, as described above. To fit *ρ*_*i*_ and *α*_*i*_, we held out one block of the random environments experiment at a time, and used the remaining 11 blocks to estimate *ρ*_*i*_, by calculating the average location of the bump of activity when the animal was at the location of each landmark on the VR track across all 11 blocks, and *α*_*i*_, by the average stability of the grid cell code around the landmark (Extended Data Fig. 5b).

Using these estimated values we simulated the dynamical model above and obtained estimated firing rates 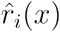 for each grid cell. We evaluated the performance of the model by computing the Pearson correlation between the estimated firing rates 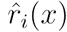 and the true firing rates *r*_*i*_(*x*). We found that for some cells the correlation was low, even when the reconstruction of the bump trajectory was quite good, indicating that our estimate of that cell’s location on the 2D sheet was poor. To avoid this issue, we additionally tried simulating grid cells at all locations on the 2D sheet, and matching true grid cells to the simulated cell with the highest Pearson correlation. We found that this procedure did not improve the model performance on cells whose locations had already been estimated well, but did improve performance on cells whose locations were estimated poorly. Finally, we found that the model’s performance tended to be higher on blocks of trials where the grid cell code was stable. To illustrate this effect, we plotted a 2D histogram of grid cell code stability (defined as the spatial correlation between the average population activity in the first half of the block and the second half of the block) and model performance (Fig. 3i).

### Position decoding analysis

To determine whether the grid cell code provided a consistent map across all three shifted environments in the tower shift task, we trained a linear decoder to predict the animal’s location on the track from grid cell population activity. Because single grid cell modules are periodic on length scales shorter than the length of the track, we trained the decoder on the combined population activity of all simultaneously recorded grid cell modules, *r*_*i*_ (*x*),*i* = 1, …,*N*_*g*_, where *N*_*g*_ represents the total number of grid cells in a recording, so that the animal’s position could be uniquely decoded. Due to the periodic nature of the VR track, we used circular-linear regression, and trained the decoder to predict two coordinates: cos(2*πx*/*L*),sin(2*πx*/*L*) using two sets of regression coefficients *β*^1^, *β*^*2*^ and a least-squares objective. The animal’s predicted position is then given by

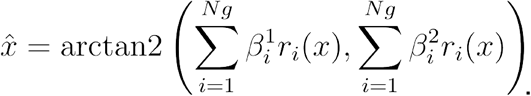

The decoder was trained on a random set of one half of the trials across all three blocks in the tower-shift environment, and evaluated on the held-out half. Cross-validated predictions, averaged across sessions and animals, are shown in Fig. 4b, right.

### Linking neural activity and behavior

For the tower-shift task, we defined the trial-by-trial stability of the grid cell code as the average spatial correlation of the grid cell population activity on a given trial with all other other trials in the block, in the 50 cm preceding the hidden reward zone. Licking error was defined as the average distance between the animals’ non-consummatory licks on that trial and the start of the hidden reward zone.

### Behavioral-timescale synaptic plasticity

To model a flexible downstream decoder which could consistently predict the hidden reward location across all three environments, we simulated a biologically plausible learning rule. The decoder was initialized with a random set of afferent weights *w*_*i*_, *i* = 1, …,*N*_*g*_ from all grid cells. Each time the animal licked and received a reward, the weights were updated in the direction of the grid cells active at the moment the animal licked.

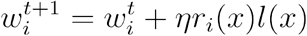

Where *l*(*x*) is an integer-valued function representing the number of consummatory licks in the spatial bin indexed by . We set *η* = 1 in our simulations. We found that the outputs

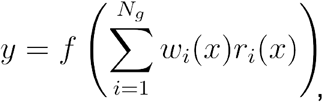

where *f* is a sigmoidal nonlinearity, rapidly adapted to deformations in the grid cell code to consistently predict hidden reward location across tower-shift environments (Fig. 4n). We simulated the effect of a plasticity knockout by allowing the decoder to adapt normally in the first environment, but setting *η* =0 in both subsequent environments, preventing the decoder from adapting to the distortions in the grid cell code induced by the tower shift. The plasticity-knockout decoder predicted that the animal would lick late both subsequent environments.

## Acknowledgements

We thank Adriana Diaz for cryosectioning and histology assistance as well as animal care and Sam Ocko for useful discussions on modeling approaches.

## Author contributions

Conceptualization (JHW, BS, SG, LMG), Methodology (JHW, BS, SG, LMG), Investigation (JHW, BS), Visualization (JHW, BS), Funding acquisition (SG, LMG), Project administration (SG, LMG), Supervision (SG, LMG), Writing – original draft (JHW, BS, LMG), Writing – review and editing (JHW, BS, SG, LMG)

## Correspondence and requests for materials

should be addressed to Lisa M. Giocomo.

## Data and code availability

Upon publication, data will be made available on a publicly accessible data repository site (e.g., figshare, DANDI, Mendeley). Custom scripts for analyzing the data will be available on GitHub.

---

**Extended Data Fig. 1:**
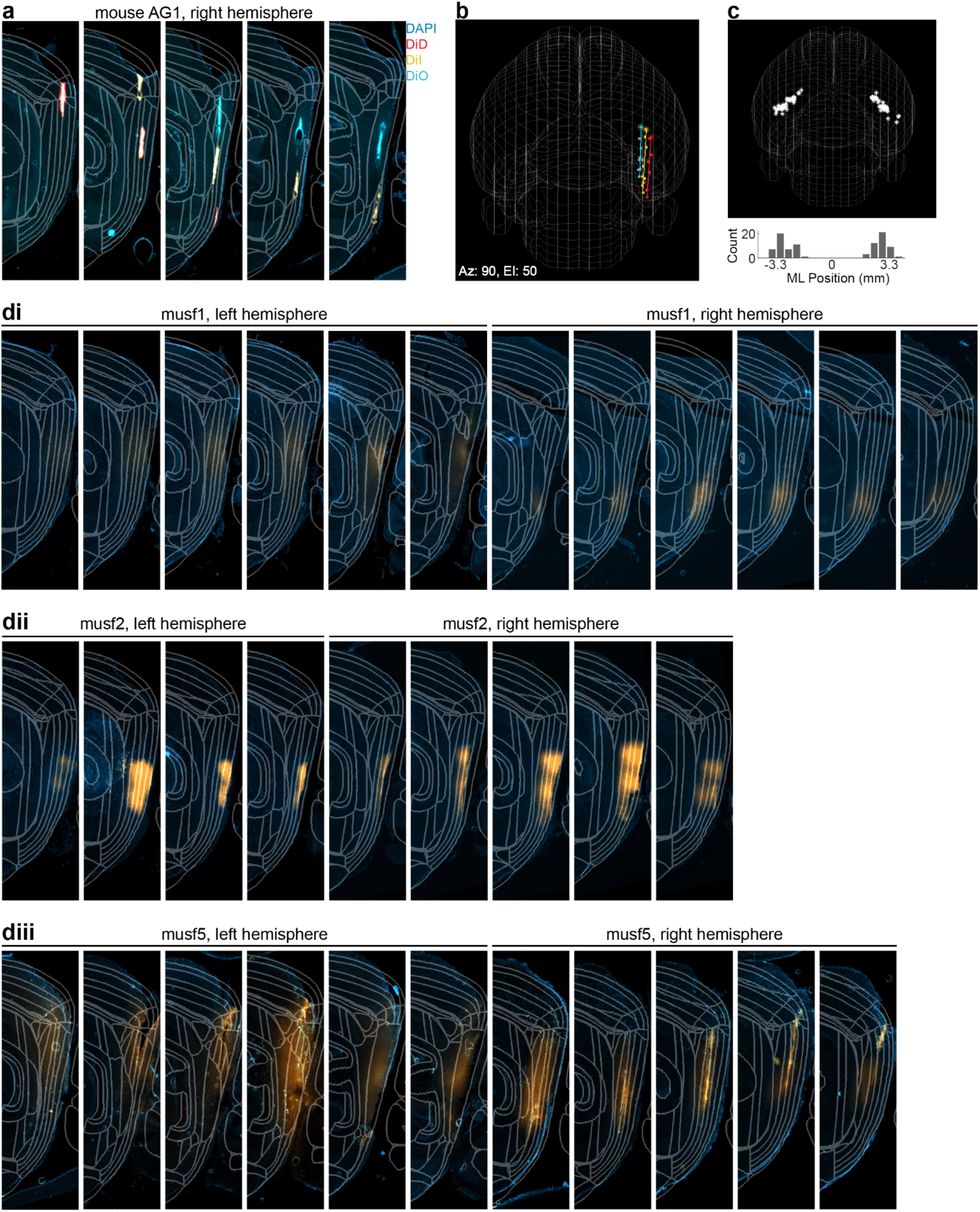
Histological confirmation of MEC targeting. **a**. Example histology from the right hemisphere of mouse AG1. Gray lines demarcate inferred boundaries between brain regions after sagittal slices were referenced to the Allen Brain Atlas using SHARP-Track 81. Slices are ordered lateral to medial from left to right. Slices were stained with DAPI, rendered in blue, to label cell nuclei. DiD, DiI, and DiO are rendered red, yellow, and turquoise, respectively. **b**. 3D reconstruction of probe locations in (a) overlaid on a mesh wire diagram of the brain. The brain is angled at azimuth 90 degrees, elevation 50 degrees for clarity of visualization. Individual colored points represented selected locations with clear dye. Solid, colored lines represent the best linear fit through the points. Starred points indicate entry points, locations where the probe first entered the brain. **c**. Quantification of entry points across all recordings (n = 93 sessions). *Top*: entry points for all recordings overlaid on a mesh wire diagram of the brain. Each star represents the entry point for an individual session. *Bottom*: histogram across all entry point medial-lateral positions relative to the midline (at 0 mm). Each bin is 400 μm. **di-diii**. Example histology showing the extent of fluorescent muscimol (in orange) spread in three animals (musf1 (di), musf2 (dii), and musf5 (diii)). Slices from the left hemisphere are ordered from most lateral to most medial, and vice versa for those from the right hemisphere. Fluorescence is primarily localized to the MEC, but a small amount can be found in the parasubiculum for a few slices

**Extended Data Fig. 2:**
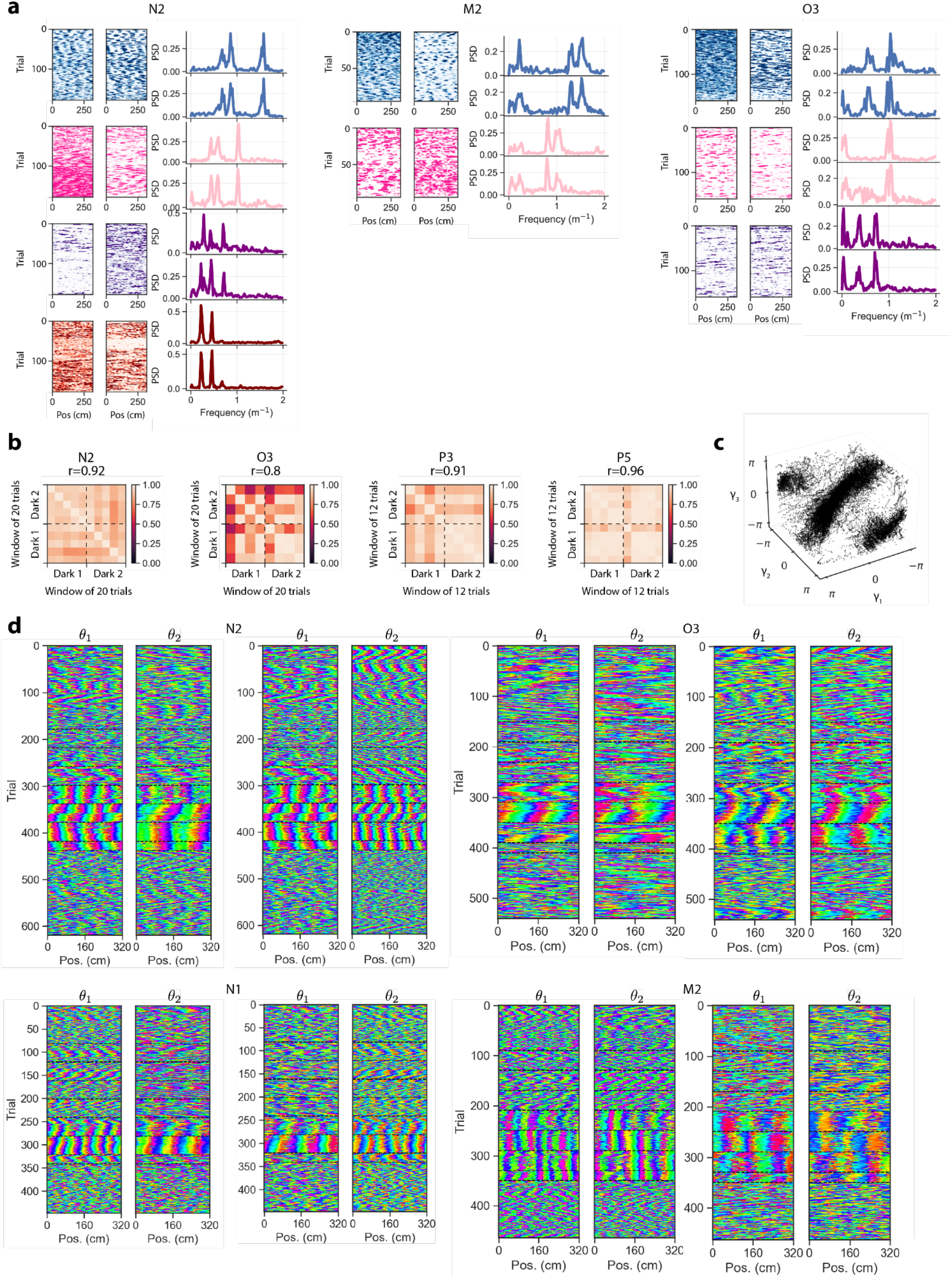
Spectral structure of the grid cell code in 1D Environments. **a**. Data from three mice shown, with mouse labels above each column. *Left:* Example spike train raster plots for grid cells from two or more different simultaneously recorded modules over 200 laps in the dark. Dots indicate spikes, color coded to match the modules identified on (*right*), sorted by overall grid scale 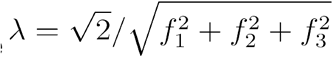 *Right:* Power spectral densities (PSDs) for each cell in (*left*), computed over a window size of 10 laps (16 meters), revealed a prominent three-peaked structure within each module. PSDs are consistent within one module, but differ between modules. **b**. Grid cell phase relationships are preserved over minutes and meters of running, over the course of an experiment. Phase relationships were defined as pairwise distances between 2D phases on the neural sheet 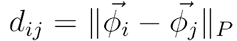. We computed the full matrix of phase relationships 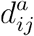 in separate windows of 10-20 trials in the dark *a*, including both windows at the beginning and end of the experiment. We then correlated the upper triangular part of 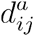 between each pair of windows *a*,*b* to obtain a matrix of window-by-window correlations. This matrix of correlations is shown in (b), for four animals and sessions. Each panel is annotated with the average pairwise correlation. **c**. Scatterplot of angle *λ*_1,2,3_ (*x*)on each of the three rings in Fig. 1j as animal (AF2) traverses the dark VR track in a single session (Methods). The three coordinates *λ*_1,2,3_ (*x*) do not take on all possible values in the 3D space, but are instead restricted to lie along a 2D subspace (module 2 *π*), indicating that they represent only two underlying degrees of freedom: the 2D coordinates of the bump of activity on the sheet. **d**. Bump trajectories, plotted as 2D coordinates on the torus: *θ*_1_ (*left*) and *θ*_2_ (*right*), for two simultaneously recorded modules from four buildup track sessions. Shown for four mice, color coded as in Fig. 2c.

**Extended Data Fig. 3:**
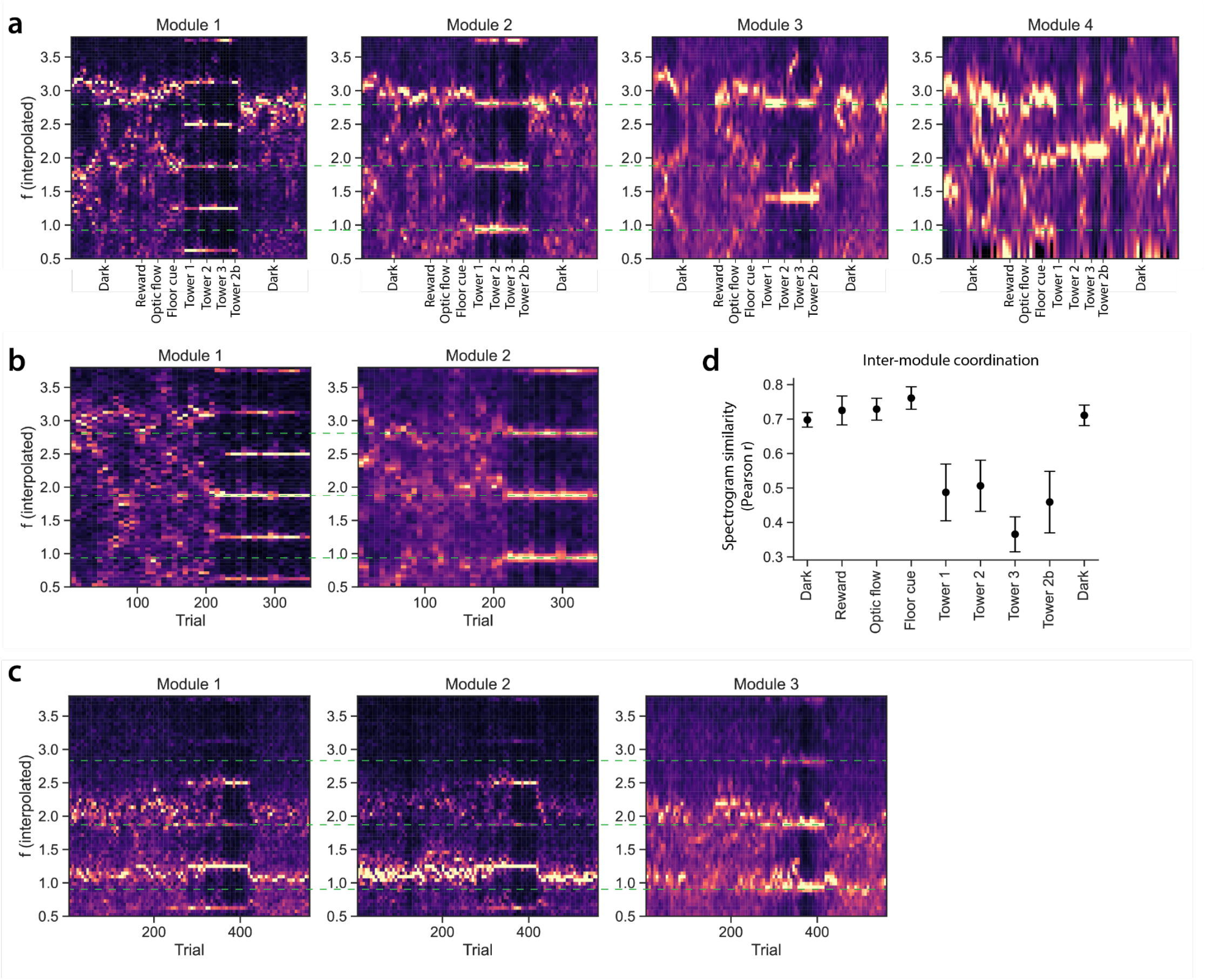
Grid cell modules drift coherently in the dark. **a-c**. Average spectrograms of grid cells from simultaneously recorded modules of increasing spatial scale, from the three sessions in Extended Data Fig. 2a. Color (dark low, white high) represents power in a given frequency *f* (y-axis). If modules rotate coherently in the dark, we would expect different modules’ spectrograms to be identical up to a stretching of the y-axis by a factor of 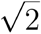 (the approximate value of the scale ratio between successive modules). To investigate this, we scale each successively larger module by a factor of 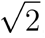, and continuously interpolate the spectrograms so that the y-axes are aligned. The interpolated spectrograms are shown in (a-c). Green lines indicate the peak frequencies for Module 2 in the presence of visual landmarks (Tower 1 - Tower 2b conditions). Plots are color coded for minimum (black) and maximum (white) values. This visualization reveals that the three peaks in the power spectrum drift coherently between modules in the dark. However, in VR, the three peaks for each module snap to incommensurate whole number frequencies. Hence the presence of landmarks disrupts the coordination between grid cell modules. We quantify this effect in panel (b). **d**. Pearson correlation between interpolated spectrograms over the course of a recording session. In the dark and in cue-poor settings, grid cell modules are tightly coordinated. This coordination is disrupted with the introduction of visual landmarks. Error bars represent standard error over windows within a block, and across the three animals and sessions (animal n = 3, session n = 3, module n = 9, cell n = 264).

**Extended Data Fig. 4:**
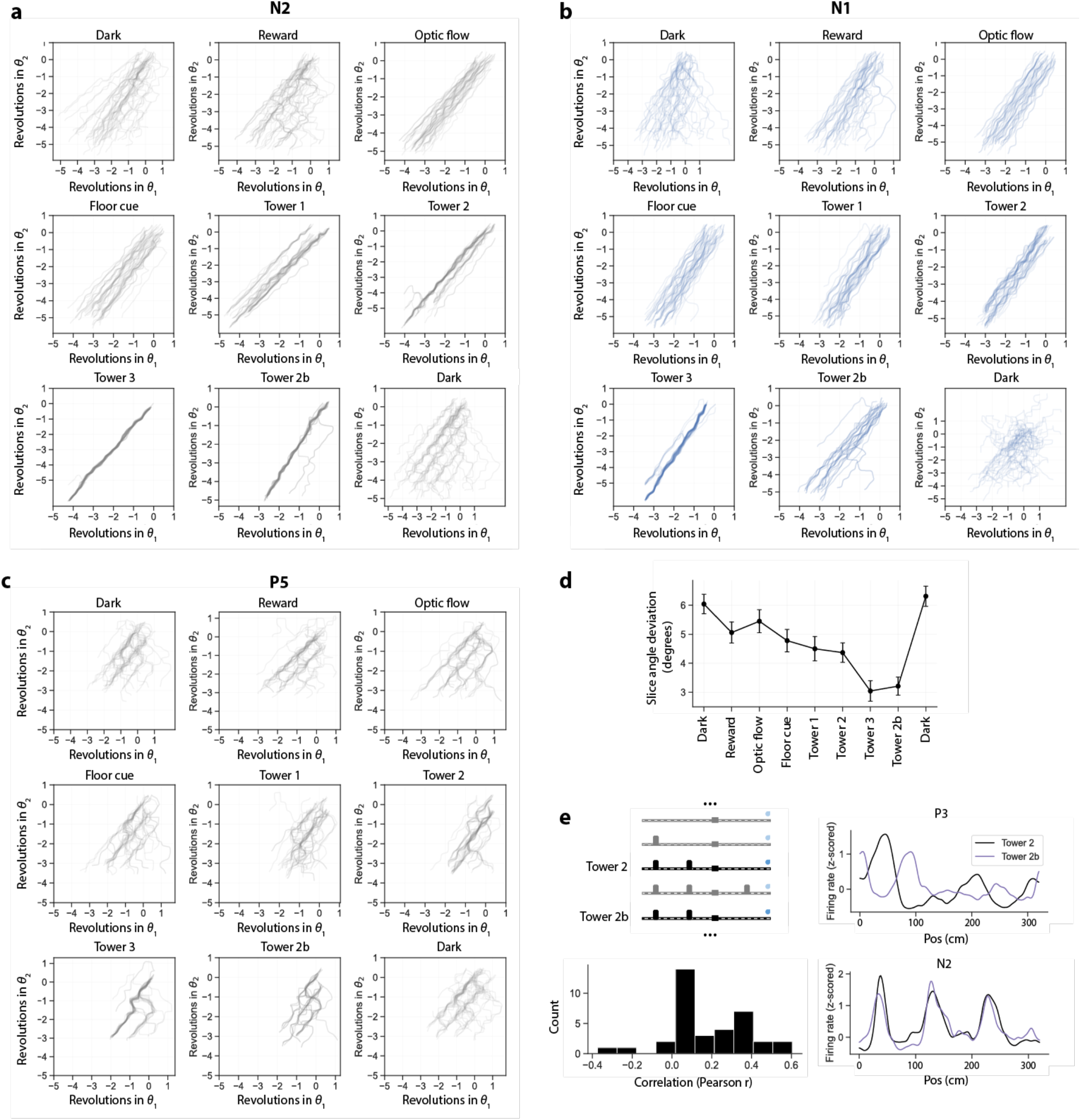
Additional activity bump trajectories and remapping statistics on the build-up track. **a-c**. Single-trial bump trajectories across all 9 blocks of the build-up track, for three different animals (Methods). **d**. Average slice angle deviation across trial blocks (circular standard deviation, session n = 14, animal n = 4, module n = 35, cell n = 747). Dots represent average deviation, and error bars standard error on the mean, across modules, sessions, and animals. **e**. Grid cell maps differ between two identical environments. *Top Left*: Schematic of the build-up track, highlighting the two visually identical environments (Tower 2 and Tower 2b blocks). *Bottom Left*: Histogram showing the median correlation of grid cell tuning curves in Tower 2 and Tower 2b blocks, across sessions and animals (animal n = 4, session n = 14, module n = 35, cell n = 747). Approximately 50% of sessions have median correlation < 0.2 between the two visually identical environments. *Top Right:* Example cell with a low correlation between Tower 2 and Tower 2b environments. *Bottom Right*: Example cell with a high correlation between Tower 2 and Tower 2b environments.

**Extended Data Fig. 5:**
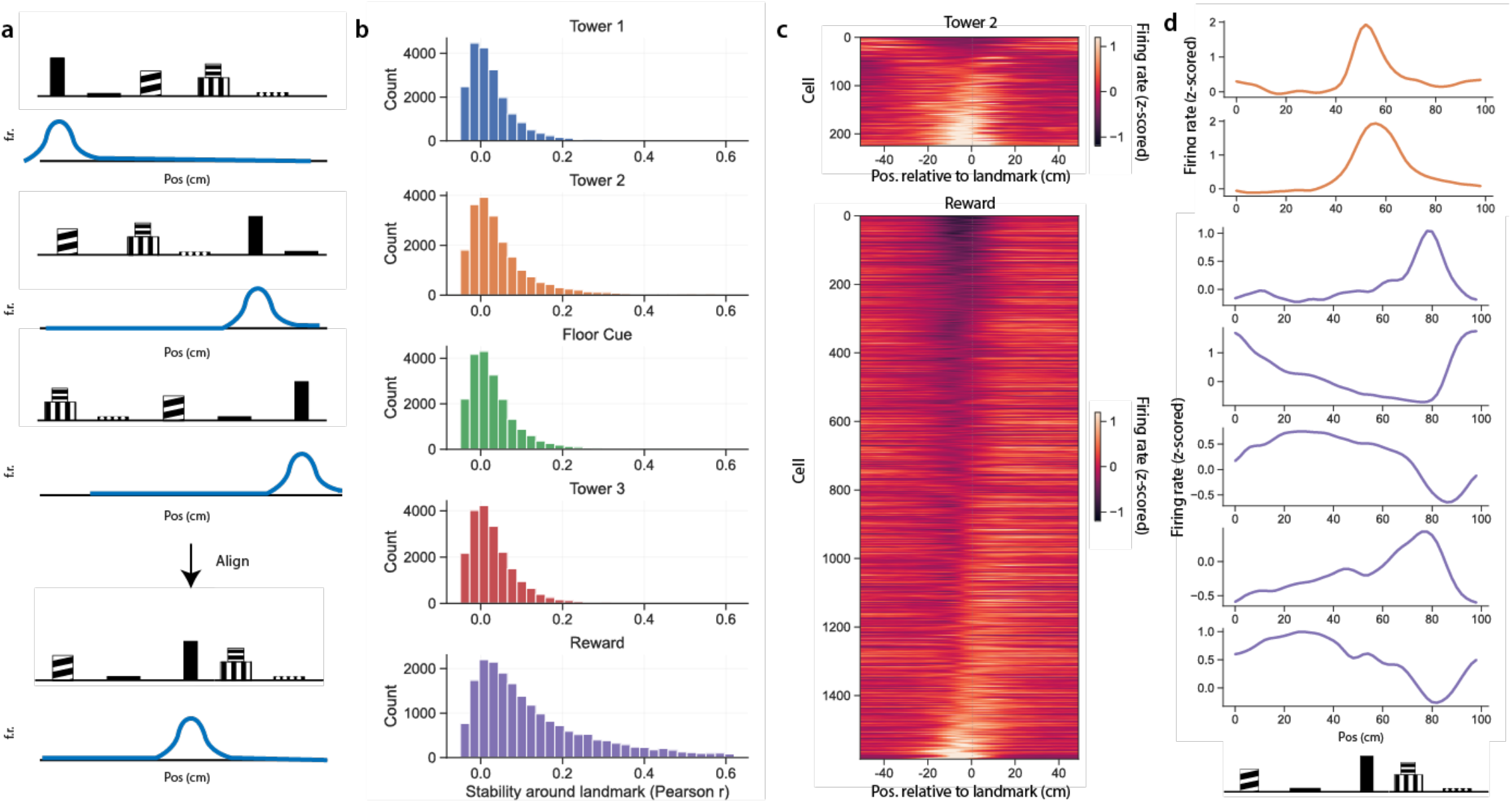
Visual response tuning in the random environment track. **a**. To determine the extent to which cells in MEC were driven by visual features, we computed tuning curves (blue line) in each block of the random environment track (as illustrated in the top three rows). We then aligned these tuning curves to each of the landmarks, one by one. A cell visually driven by landmark A, when aligned to landmark A, should have a similar tuning curve across environments (as illustrated by the bottom row). **b**. Quantification of the similarity of neural activity for each landmark, analyzed as described in (a). Each graph represents the stability of neurons (n = 68,484) with respect to each landmark. Stability is defined as the average spatial correlation of landmark-aligned maps across all 12 random environment blocks. Note there were very few cells with a high similarity (that is, with similar responses to the same visual landmark across environments). The reward had the strongest “pinning”, with the greatest number of cells exhibiting consistent tuning to the reward. Of the visual landmarks, Tower 2 had the largest number of tuned cells. **c**. Heatmap of tuning curves of all cells considered “stable” (Pearson r > 0.4) for Tower 2 (orange) and for the reward (purple), averaged across random environments. **d**. Example aligned tuning curves for individual cells, for Tower 2 and for the reward, averaged across random environments.

**Extended Data Fig. 6:**
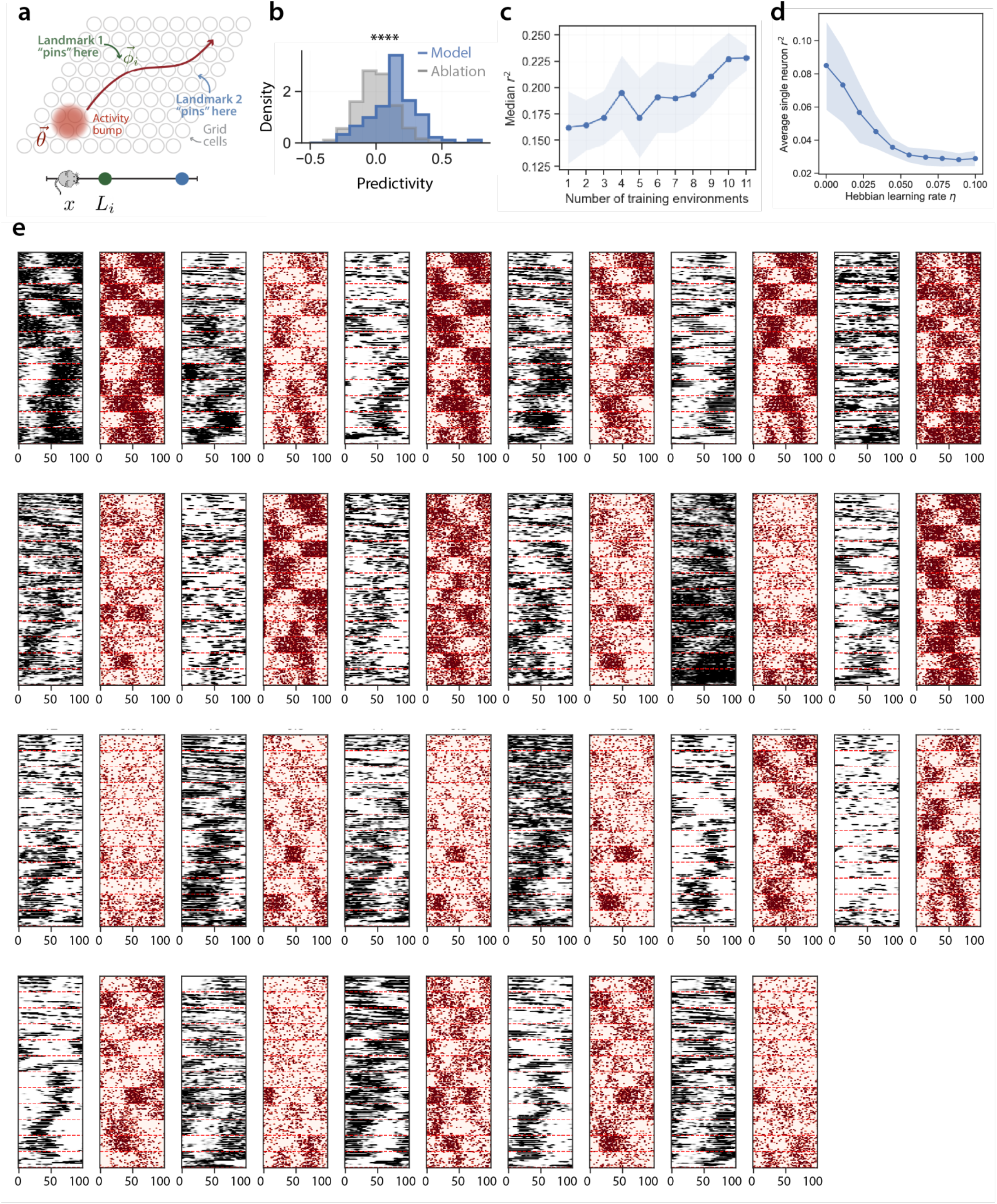
Model details and performance. **a**. Schematic of 2D dynamical model of grid cell population code. **b**. In addition to comparing to a random shuffle in Fig. 3h, we performed an ablation to confirm that the model uses the spatial locations of the landmarks to predict the grid cells’ activity patterns in a novel environment, and does not simply reproduce their average firing properties. To verify this, we fed randomly shuffled landmark locations, rather than the true landmark locations, to the model in each held-out environment. This ablation disrupted the performance of the model to no better than a random shuffle (Fig 3h), indicating that knowledge of the landmark locations in the novel environment is crucial to predict grid cells’ activity patterns. **c**. Model performance improves as we increase the number of random environments used to estimate the model parameters (“pinning phases”, Methods). **d**. Neural activity is better explained by a fixed, non-plastic circuit than a plastic circuit where pinning phases *ρi* evolve from trial to trial to minimize their distance to the bump of activity when the animal is near the corresponding landmark, with learning rate *η* (see 21 for details on this Hebbian learning rule). **f**. True grid cell rasters (black) and simulated grid cell responses for all cells within a single session (red).

**Extended Data Fig. 7:**
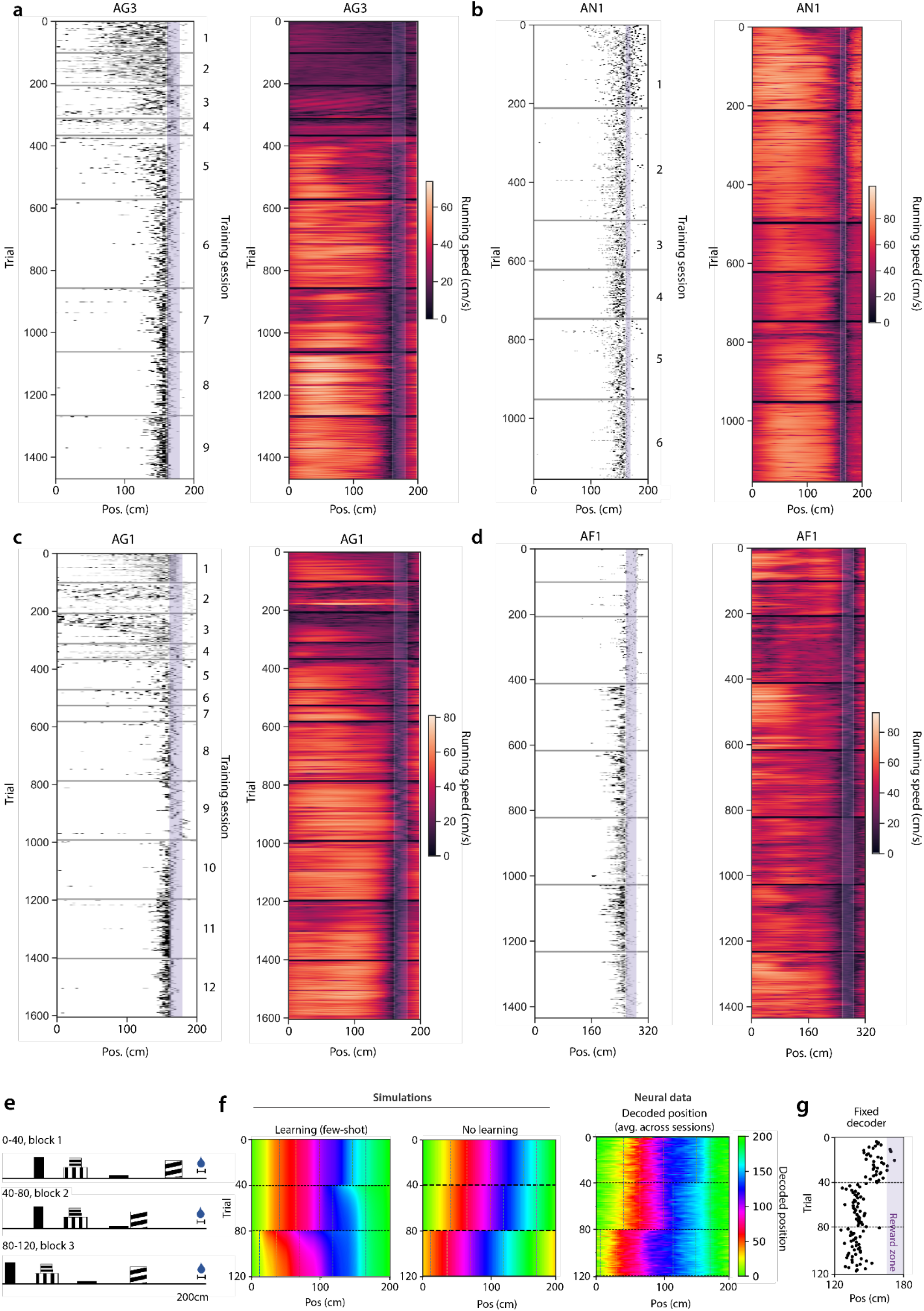
Animal performance throughout training on the hidden reward task and additional decoder simulations. **a-d**. Behavioral performance on the hidden reward task over the course of training for four mice. *Left*: Non-consummatory licks, dots indicate licks and the purple bar indicates reward zone. *Right*: Running speed. Horizontal lines indicate training sessions. Over the course of training, animals learn to slow down and lick in anticipation of the hidden reward zone (purple). For training details see Methods. **e-g**. Neural activity in the “backward shift condition”, analogous to what is shown in Fig. 4a, b, d. Behavior is shown in Fig. 4g. **e**. Schematic of the hidden reward task for the “backward shift condition”. The unmarked reward zone is indicated by the water drop. The mouse traveled a variable distance from the start of each trial to the first landmark. **f**. *Left-Middle*: Decoded position from a population of simulated grid cells which exhibit few-shot learning (*left*) and no learning (*middle*) on the “backward shift condition”. Horizontal black dotted lines correspond to the trial in which the location of the visual cue shifts. Vertical lines indicate the location of the landmarks for each block of trials. *Right*: Decoded spatial position from a linear decoder trained on the population activity of simultaneously recorded grid cells in a tower-shift experiment, averaged over modules, sessions, and animals (animal n = 5, session n = 10, module n = 29, cell = 1325). **g**. Predicted lick locations for a fixed decoder trained on all three blocks in the “backward shift condition”. Dots represent licks.

**Extended Data Fig. 8:**
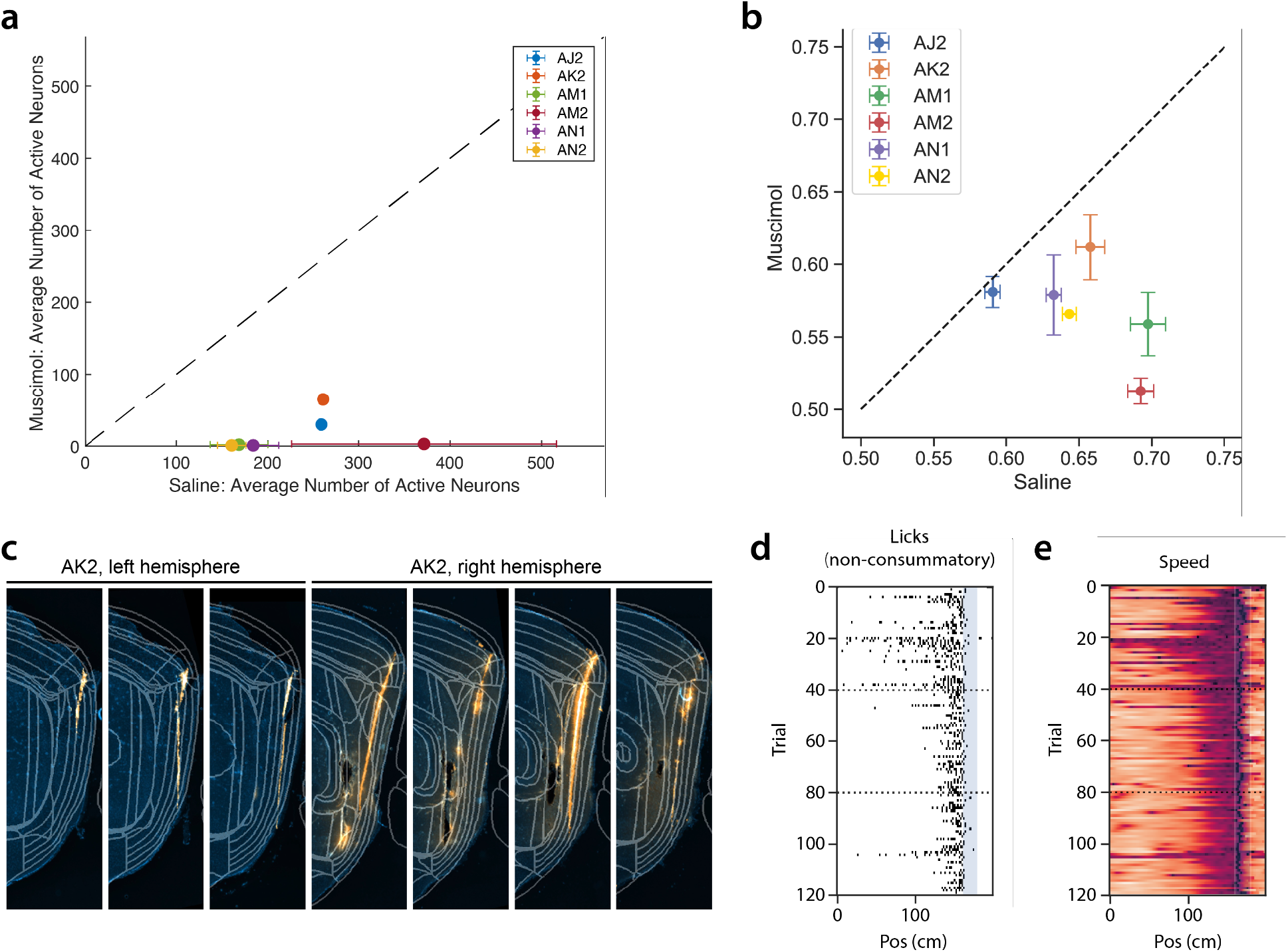
Muscimol-induced disruption of neural activity is correlated with navigational impairments. **a**. Scatterplot of the average number of well-isolated, active neurons recorded in MEC following bilateral infusion of saline or muscimol. Each dot represents the mean for a single animal, with standard error of mean error bars where appropriate (> 1 recording for a given condition). Dashed black line represents the unity line. Pooled across all sessions, muscimol significantly reduced the number of active units (two-tailed unpaired t-test, p < 1e-4, animal n = 6, saline sessions n = 10, muscimol sessions n = 8). Note that animal AK2, represented by the orange point, is a potential outlier, since the number of recorded active units following muscimol infusion was greater than those for other animals. See panels (c-e) for related histology and behavior from the muscimol session for this animal. **b**. Scatterplot of average grid cell stability (quantified by average Pearson r correlation of grid cell population activity on one trial to other trials within a block) following bilateral infusion of saline or muscimol. Each dot represents the average Pearson r for a single animal with standard error of mean error bars where appropriate (> 1 recording for a given condition). Dashed black line represents the unity line. Compared to saline, muscimol significantly reduced grid map stability (Wilcoxon rank sum test, p < 1e-4, animal n = 6, saline sessions n = 10, muscimol sessions n = 8). **c**. Histology from animal AK2 showing incomplete spread of fluorescent muscimol in the left hemisphere of MEC, consistent with the higher number of concurrently recorded units from the same hemisphere. **d-e**. Animal AK2’s licking (d) and running (e) behavior on the hidden reward track following bilateral fluorescent muscimol injection. Consistent with the incomplete silencing of MEC activity in (a) and partial spread of fluorescent muscimol in (c), licking and running behavior were not as impaired as for other animals with more complete MEC silencing.

**Extended Data Table 1:** Tabulation of all of the animals and sessions they ran in this manuscript. The top half of the table enumerates the number of sessions a given animal contributed for each experiment. The bottom half of the table summarizes the total number of animals and sessions for each experiment.

| Animal | # Build-Up Track | # of Random Environment | # of Hidden Reward | # of Hidden Reward with Muscimol and Neuropixels Recording | # of Hidden Reward with Muscimol and no Neuropixels Recording | # of Hidden Reward with Fluorescent Muscimol and Neuropixels Recording | # of Hidden Reward with Fluorescent Muscimol and no Neuropixels Recording | # of Hidden Reward with Saline and Neuropixels Recording | # of Hidden Reward with Saline and no Neuropixels Recording | # of Fluorescent Muscimol only |
| --- | --- | --- | --- | --- | --- | --- | --- | --- | --- | --- |
| M2 | 6 | 0 | 0 | 0 | 0 | 0 | 0 | 0 | 0 | 0 |
| N1 | 5 | 0 | 0 | 0 | 0 | 0 | 0 | 0 | 0 | 0 |
| N2 | 5 | 0 | 0 | 0 | 0 | 0 | 0 | 0 | 0 | 0 |
| P3 | 3 | 0 | 0 | 0 | 0 | 0 | 0 | 0 | 0 | 0 |
| AF3 | 0 | 3 | 0 | 0 | 0 | 0 | 0 | 0 | 0 | 0 |
| AF4 | 0 | 4 | 0 | 0 | 0 | 0 | 0 | 0 | 0 | 0 |
| AG1 | 0 | 3 | 6 | 0 | 0 | 0 | 0 | 0 | 0 | 0 |
| AJ2 | 0 | 3 | 0 | 1 | 1 | 0 | 0 | 1 | 2 | 0 |
| AK1 | 0 | 3 | 1 | 0 | 0 | 0 | 1 | 1 | 0 | 0 |
| AK2 | 0 | 2 | 1 | 0 | 0 | 1 | 0 | 1 | 0 | 0 |
| AL1 | 0 | 2 | 0 | 0 | 0 | 0 | 0 | 0 | 0 | 0 |
| AM1 | 0 | 4 | 3 | 2 | 0 | 0 | 0 | 2 | 0 | 0 |
| AM2 | 0 | 6 | 2 | 1 | 0 | 0 | 0 | 2 | 0 | 0 |
| AN1 | 0 | 4 | 1 | 1 | 0 | 0 | 0 | 2 | 0 | 0 |
| AN2 | 0 | 5 | 2 | 2 | 0 | 0 | 0 | 2 | 0 | 0 |
| musf1 | 0 | 0 | 0 | 0 | 0 | 0 | 0 | 0 | 0 | 1 |
| musf2 | 0 | 0 | 0 | 0 | 0 | 0 | 0 | 0 | 0 | 1 |
| musf5 | 0 | 0 | 0 | 0 | 0 | 0 | 0 | 0 | 0 | 1 |
| musf6 | 0 | 0 | 0 | 0 | 0 | 0 | 0 | 0 | 0 | 1 |
|  | # of build-up track sessions | # of random environment sessions | # of hidden reward sessions | # of hidden reward sessions with muscimol and neuropixels | # of hidden reward sessions with muscimol and no neuropixels | # of hidden reward sessions with fluorescent muscimol and neuropixels | # of hidden reward sessions with fluorescent muscimol and no neuropixels | # of hidden reward sessions with saline and neuropixels | # of hidden reward sessions with saline and no neuropixels |  |
|  | 19 | 39 | 16 | 7 | 1 | 1 | 1 | 11 | 2 |  |
|  | # of animals build-up track | # of animals random environment | # of animals hidden reward | # of animals hidden reward with muscimol and neuropixels | # of animals hidden reward with muscimol and no neuropixels | # of animals hidden reward with fluorescent muscimol and neuropixels | # of animals hidden reward with fluorescent muscimol and no neuropixels | # of animals hidden reward with saline and neuropixels | # of animals hidden reward with saline and no neuropixels | # animals with fluorescent muscimol |
|  | 4 | 11 | 7 | 5 | 1 | 1 | 1 | 7 | 1 | 6 |

